# Primate-specific response of astrocytes to stroke limits peripheral macrophage infiltration

**DOI:** 10.1101/2020.05.08.083501

**Authors:** Anthony G. Boghdadi, Joshua Spurrier, Leon Teo, Mingfeng Li, Mario Skarica, Benjamin Cao, William Kwan, Tobias D. Merson, Susie K. Nilsson, Nenad Sestan, Stephen M. Strittmatter, James A. Bourne

## Abstract

Reactive astrocytes play critical roles after brain injuries but their precise function in stroke is not well defined. Here, we utilized single nuclei transcriptomics to characterize astrocytes after ischemic stroke in nonhuman primate (NHP) marmoset monkey primary visual cortex. We identified 19 putative subtypes of astrocytes from injured and uninjured brain hemispheres and observed nearly complete segregation between stroke and control astrocyte clusters. We then screened for genes that might be limiting stroke recovery and discovered that one neurite-outgrowth inhibitory protein, NogoA, previously associated with oligodendrocytes but not astrocytes, was expressed in numerous reactive astrocyte subtypes. NogoA upregulation on reactive astrocytes was confirmed *in vivo* for NHP and human, but not observed to the same extent in rodent. Further *in vivo* and *in vitro* studies determined that NogoA mediated an anti-inflammatory response which limits deeper infiltration of peripheral macrophages from the lesion during the subacute post-stroke period. Specifically, these findings are relevant to the development of NogoA-targeting therapies shortly after ischemic stroke. Our findings have uncovered the complexity and species specificity of astrocyte responses, which need to be considered more when investigating novel therapeutics for brain injury.

---

While thrombolytic therapies have proven beneficial for ischemic stroke recovery, decades of effort to translate neuroprotective discoveries from rodent experiments to the clinical environ have been typified by numerous failures^1^. There are likely multiple explanations, but a focus on rodents for discovery research and the preeminent focus on neuronal responses may be crucial factors. For this reason, we investigated astroglial responses in a nonhuman primate (NHP) model of ischemic stroke, which has previously been demonstrated to be more akin to the human condition^2^.

It is well established that astrocytes are crucial players in the pathogenesis of brain injuries, such as ischemic stroke, yet the primary focus has remained on the neuron^3^. In response to exogenous or endogenous stimuli arising from injury, disease or aging, astrocytes are triggered into their reactive state (reactive astrocytes), characterized by drastic changes to their gene expression, morphology and function. Although it is recognized that reactive astrocytes can exacerbate secondary neurodegeneration, recent studies have highlighted that they are a highly heterogeneous population of cells that can also provide crucial neuroprotection for surviving neurons in response to CNS pathologies^4^. Recent transcriptome analyses have defined distinct subtypes of reactive astrocytes with neurotoxic^5^ or neuroprotective^6^ properties that are induced after specific types of CNS injury in mice, highlighting the injury-dependent plasticity of reactive astrocytes after brain injury. Here, we provide the first catalogue of NHP astrocyte diversity in a clinically translatable stroke model and identify putative therapeutic targets from the reactive astrocyte-mediated pathogenesis of ischemic stroke.

Single-nuclei RNA sequencing (snRNAseq) identified 19 discrete subtypes of astrocytes, revealing marked diversity and activation in marmoset. Unexpectedly, we observed that a neurite-outgrowth inhibiting protein associated with myelinating oligodendrocytes (NogoA)^7, 8^ is expressed by certain reactive astrocytes. On the one hand, this primate-specific response provides a local secondary source of neurite outgrowth inhibition, potentially reinforcing oligodendrocyte effects. In addition, we present evidence that a collar of reactive astrocytes expressing NogoA play an anti-inflammatory role in the subacute stroke period to limit peripheral macrophage infiltration from the ischemic zone (adjacent to the ischemic core). Therefore, immediate post-stroke blockade of NogoA action may exacerbate inflammation selectively in primate species, including humans. These data highlight the pronounced diversity of astrocyte populations and their responses to ischemic damage, as well as the critical role of primate-specific CNS responses to injury.

## Results

### Single-nuclei RNAseq revealed 19 unique astrocyte subtypes in the primate post-ischemic stroke

We used single-nuclei RNAseq (snRNAseq) on the 10X platform to profile NHP astrocytes following ischemic stroke. Adult marmosets (n=3) received unilateral injections of endothelin-1 (ET-1) to the primary visual cortex (V1), with tissues collected 7 days later^2^. The contralateral hemisphere was used as an internal control (**Fig. 1a; Supplementary Fig. 1a**). Single nuclei (134,861 total; **Supplementary Fig. 1c)** were isolated from snap frozen tissue. Following sequencing, individual nuclei underwent unsupervised clustering and were arranged by uniform manifold approximation and projection (UMAP) for visualization, with nuclei from the grey matter (GM) and white matter (WM) of V1 being analyzed separately (**Fig. 1b**). Clusters were manually assigned to neural cell types based on the expression patterns of known cell-type-specific markers (**Fig. 1c, Supplementary Fig. 1d, Supplementary Table 1**).

**Figure 1.**
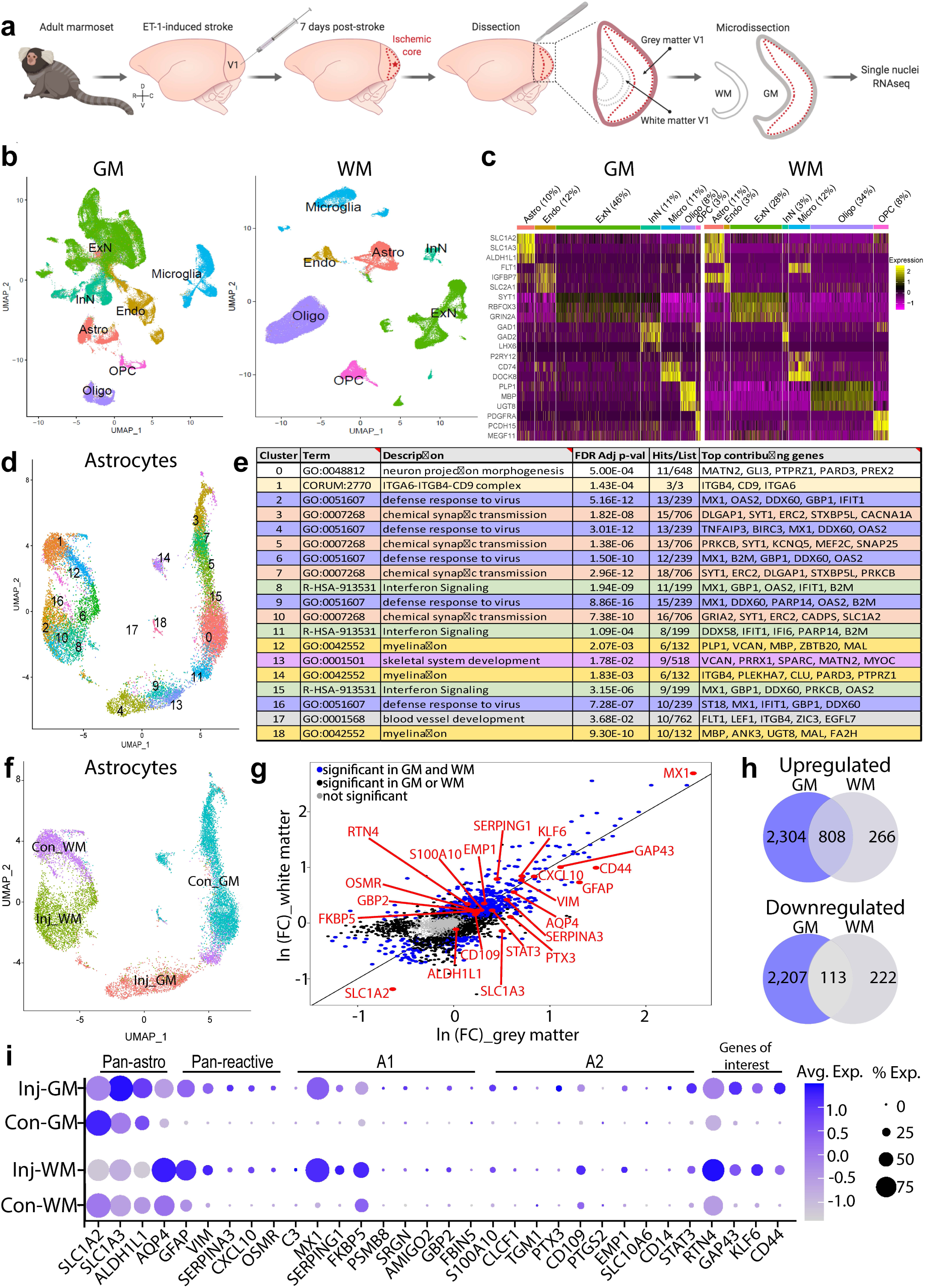
Molecular classification of marmoset post-ischemic stroke reactive astrocytes **(a)** Schematic depicting endothelin-1 (ET-1)-induced ischemic stroke in adult marmoset primary visual cortex (V1) and tissues isolated for single-nuclei 10x genomic sequencing. The injury was induced in the left hemisphere (Injured; Inj); the contralateral (Con) hemisphere was used as a control for transcriptomic analyses. Grey matter (GM) and white matter (WM) were collected and isolated from both hemispheres. **(b)** UMAP visualization of single nuclei from GM or WM, colored by cell type. **(c)** Heat map colored by single nuclei gene expression of cell-type specific markers. Heat map of additional markers used for cell-type assignment is in **Supplementary Fig. 1d. (d)** UMAP visualization of single astrocyte nuclei from GM and WM samples, colored by cluster; clusters are numbered based on the size of the cluster. **(e)** Metascape gene ontology analysis using the top 50 genes from each astrocyte subtype; identical GO processes are color-coded. Top GO process is presented here; full GO results are found in **Supplementary Table 3. (f)** UMAP visualization of single astrocyte nuclei, colored by injury status and region. Inj: injured/ stroke hemisphere; Con: contralateral hemisphere. **(g)** Differentially expressed genes (DEGs) between injured and contralateral hemispheres; the natural log of the fold change from either region is plotted on the axes. Genes were deemed significant if they exhibited >0.25 log(e)-transformed fold change in either direction, or if the ratio of nuclei expressing the gene between Injured and Contralateral nuclei was >1.25 and if they were expressed in >10% of nuclei in both groups. Black dots represent significant DEGs in either GM or WM, while blue dots represent DEGs that are significantly altered in both GM and WM. Grey dots are genes with unaltered expression in both regions. Genes labeled in red are previously described astrocyte markers or particular genes of interest. Specific fold-change values and p-values are in **Supplementary Fig. 1f. (h)** Summary of upregulated and downregulated DEGs in GM and WM. **(i)** Average **e**xpression of pan-astrocyte and reactive astrocyte markers by injury status and region. The size of the dots indicates the percent of nuclei within that group that are expressing the gene at any level, while color indicates average expression from all nuclei within that group. “A1” and “A2” refer to markers of neurotoxic and neuroprotective astrocytes, respectively, as previously described (Liddelow et al, 2017). “Genes of interest” include NogoA (RTN4) and a subset of genes with concordant expression: growth associated protein 43 (GAP43), Kruppel-like factor 6 (KLG6), and cell-surface glycoprotein-44 (CD44).

As there has already been considerable focus on neuronal changes following ischemic injury, we sought to profile the changes to astrocytes after stroke. Given that different astrocyte subtypes populate the WM and GM^9^, astrocyte nuclei from the two regions from both injured and contralateral hemispheres were first isolated before being pooled together and subclustered using the Seurat algorithm. We identified a total of 19 astrocyte subtypes (**Fig. 1d**), each enriched for specific ontology processes (**Fig. 1e**). Three distinct phenotypes of reactive astrocytes have previously been described in rodents: neurotoxic “A1” astrocytes ^5^; neuroprotective “A2” astrocytes ^6^; and, “scar-forming” astrocytes ^10^. Analysis of the 19 marmoset astrocyte subtypes revealed that none of the astrocytes strongly upregulated the full panel of either A1 or A2 markers (**Supplementary Fig. 2**). Notably, certain subtypes are marked by genes not typically expressed in astrocytes, such as myelin basic protein (MBP), a structural component of myelin expressed by oligodendrocytes (**Supplementary Table 1**). These subtypes could signify intermediate states of astrocytes, such as polydendrocytes^11^. Together these data suggest that there are many more subtypes of reactive astrocytes following stroke than previously described.

To further determine specific injury-induced changes, nuclei were grouped based on injury status and region (Injury-GM, Contralateral-GM, Injury-WM, Contralateral-WM; **Fig. 1f**). Post-stroke changes are observed in both GM and WM, as subclustering of the astrocyte nuclei reveals near-total separation in UMAP space between the injured and contralateral samples. Injured samples were compared directly to the respective contralateral (control) to identify differentially expressed genes (5,432 in GM, and 1,409 in WM; **Fig. 1g, Supplementary Table 2**). More genes were affected in GM than in WM, and the majority of genes were upregulated, particularly in the subset of genes common between both regions (**Fig. 1h**). Astrocytes from both Inj-GM and Inj-WM exhibited differential expression of: multiple markers of reactive astrocytes (GFAP, VIM, SERPINA3, CXCL10); immunomodulatory genes, including genes involved in defense response to virus (IFI6, GBP1, IFIT1, IFIT3, DDX58); and, genes contributing to various cytokine/interferon signaling pathways (B2M, STAT1, IRF1, NLRC5) (**Fig. 1g, i; Supplementary Tables 2-3**). The transcriptional activator Klf-6, previously reported to be upregulated following MCAO^6^, was also found to be significantly upregulated in both Inj-GM and Inj-WM (**Fig. 1g, i**). Similarly, CD44, the major surface hyaluronan (HA) receptor, which plays a role in cell-matrix adhesion and cell migration^12^, was also upregulated in both injured regions. A number of neuroprotective genes were also identified. Astrocytic expression of growth associated protein 43 (GAP43) was recently shown to mediate glial plasticity and promote neuroprotection^13^; here, GAP43 was significantly upregulated in both Inj-GM and Inj-WM astrocytes (**Fig. 1g,i**). Taken together these data indicate that primate astrocytes are involved in a variety of immunomodulatory and neuroprotective functions after stroke but are different from the classical A1 ‘neuroprotective’ subtype identified in rodents.

Notably, it was observed that NogoA, previously described as an oligodendrocyte marker^14^ and known to inhibit neurite outgrowth^15, 16^, was significantly upregulated in astrocytes from both Inj-GM and Inj-WM tissue (**Fig. 1g,i; Fig. 2a**). Expression of NogoA, along with GAP43, KLF-6, and CD44, in individual nuclei revealed a robust distinction between injury and contralateral samples; both the percent of nuclei expressing these genes and the overall level of expression were increased (**Fig. 2a**). These findings were validated by immunohistochemistry in tissues from control and one-week post-stroke marmosets, where we observed increases in the number of astrocytes colocalizing NogoA and GAP43, KLF-6, or CD44 immunoreactivities (**Fig. 2b-c**; **Supplementary Fig. 3**). These data demonstrate that our transcriptomic results are consistent with tissue protein levels.

**Figure 2.**
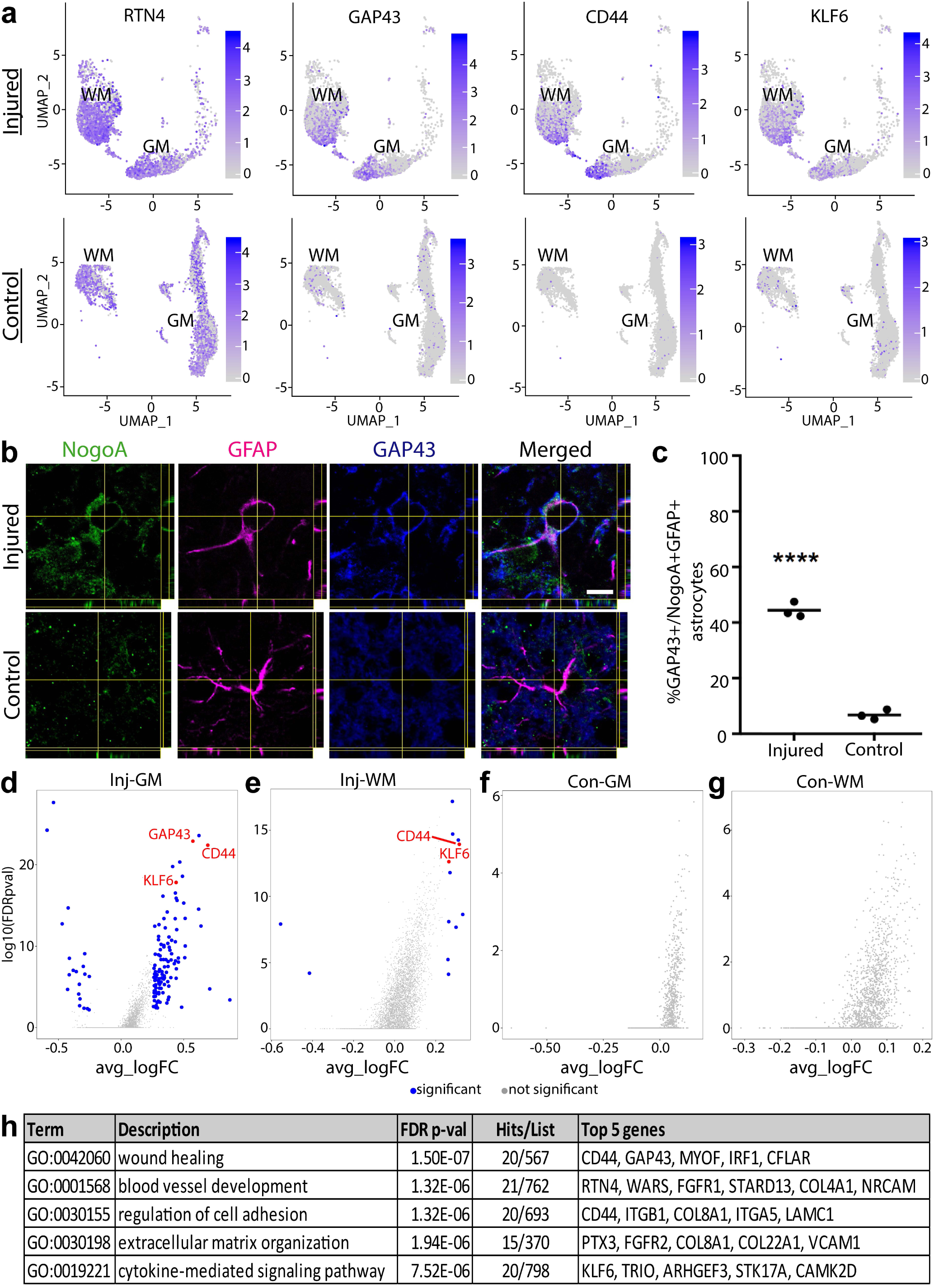
Characterization of marmoset post-ischemic stroke NogoA-positive reactive astrocytes. **(a)** Nuclei-specific expression of NogoA (RTN4), growth associated protein 43 (GAP43), cell-surface glycoprotein-44 (CD44), and Kruppel-like factor 6 (KLG6). **(b)** Immunohistochemical validation of GAP43 upregulation in NogoA-positive reactive astrocytes from a separate cohort of adult marmosets. Injured = 7 days post-injury (ET-1 injection); Control = non-injected marmoset. Scale bars: 10 µm. **(c)** Quantification of GAP43+/NogoA+/GFAP+ triple positive astrocytes from **(b)**. Each dot represents the mean percentage for individual animals where at least 50 cells were counted over a minimum of 3 sections at high magnification with stacks to confirm colocalization of NogoA, GFAP, and GAP43. Statistical significance was determined using an unpaired t-test. **(d-g)** Differentially expressed genes between NogoA-positive and NogoA-negative nuclei in **(d)** injured GM, **(e)** injured WM, **(f)** contralateral GM, and **(g)** contralateral WM. **(h)** Metascape gene ontology analysis using all significant genes from **(d)**. Using all significant hits from **(e)** did not result in any significant enrichment of ontological processes, and no significant genes were identified in **(f)** or **(g)**.

To probe potential mechanisms associated with NogoA, we examined differentially expressed genes between NogoA-positive and NogoA-negative nuclei in each of the four groups. 130 genes were identified in Inj-GM (110 upregulated in concert with NogoA and 20 downregulated) (**Fig. 2d**), while only 13 (11 co-upregulated and 2 downregulated) were observed in Inj-WM (**Fig. 2e**). No genes were differentially expressed between NogoA positive and negative nuclei in either of the contralateral groups (**Fig. 2f-g**). Ontology analysis revealed that the gene set correlating with NogoA expression in Inj-GM is enriched for wound healing, regulation of cell adhesion, extracellular matrix organization, and cytokine-mediated signaling pathway, among others (**Fig. 2h, Supplementary Table 3**). These data indicate that NogoA may be involved in functions other than those it is typically associated with in the literature, including, but not limited to immunomodulatory reactive astrocyte functions.

### Reactive astrocytes express neurite-outgrowth inhibitory protein NogoA in primates post-ischemic stroke

The best characterized sequela of NogoA expression after CNS injuries is inhibition of neurite outgrowth^7, 8^, through neuron-oligodendrocyte, axon-interactions. Having identified that reactive astrocytes express NogoA in the marmoset one-week post-stroke (**Fig. 1g,i; 2a-c**), and given this had not been previously described in rodent literature, we sought to investigate astrocytic NogoA expression at the equivalent post-stroke time-point of 3 and 7 DPI in mouse and marmoset, respectively. Our analysis revealed fewer than 50% NogoA+ astrocytes adjacent to the ischemic core in mouse cortex (**Fig. 3a, c**), with minimal and punctate labeling compared to the widespread and uniform NogoA expression observed in almost all marmoset astrocytes at 7 DPI within the ischemic zone (**Fig. 3b, c**). Furthermore, the fluorescence intensity of NogoA immunoreactivity in astrocytes of the mouse was negligible relative to that in the marmoset (**Fig. 3d**). Together, these data indicated that there is a marked difference between mouse and marmoset in terms of upregulation of NogoA on reactive astrocytes post-stroke.

**Figure 3.**
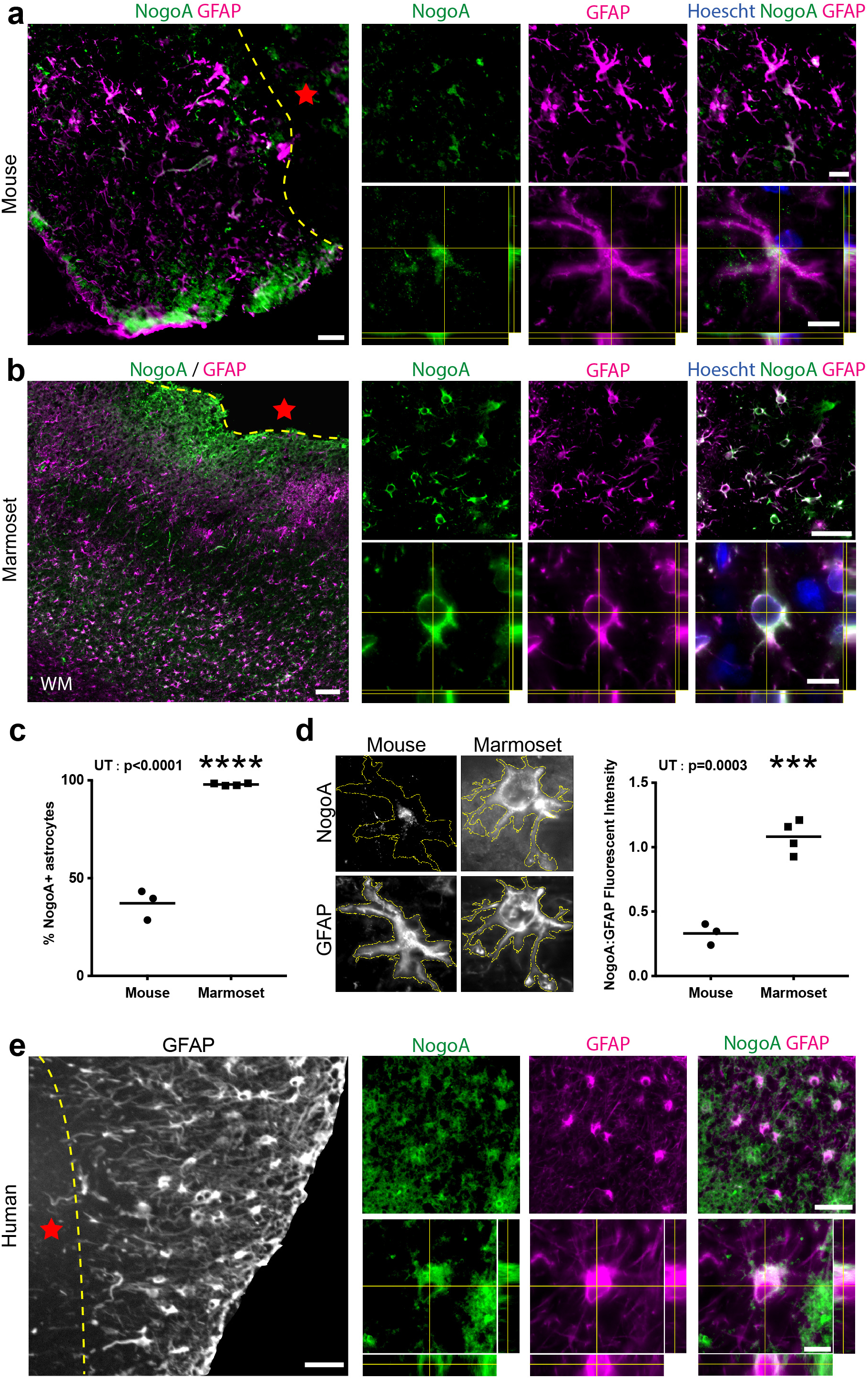
NogoA expression after ischemic stroke in mouse, marmoset and human. (**a, b, e**) Representative immunofluorescent photomicrographs and stacks with orthogonal views showing NogoA colocalization with GFAP+ cells in mouse (**a**) and primate (**b**: marmoset; **e**: human) cortical tissue adjacent to the ischemic core 3 and 7 days post-ischemic stroke, respectively. (**c**) Quantification of NogoA+/ GFAP+ cells in mouse versus marmoset expressed as a percentage of total counted astrocytes. Each dot represents the mean percentage for individual animals, where at least 50 cells were counted over a minimum of 3 sections at high magnification with stacks to confirm colocalization of NogoA and GFAP (**d**) Quantification of NogoA fluorescent intensity, normalized against GFAP, in mouse versus marmoset. Each dot represents the mean fluorescent intensity of NogoA:GFAP for individual animals where at least 10 cells from previous counts were analyzed at high magnification. NogoA: neurite outgrowth inhibitor A; GFAP: glial fibrillary acidic protein, hoechst: nuclear stain; WM: white matter; yellow dotted line: border of ischemic core; red star: ischemic core; scale bars: 100 µm (**a, b, e**: left), 50µm (**a, b, e**: top), 10 µm (**a, b, e**: bottom); statistical test: unpaired T test (UT); ***: p<0.001; ****: p<0.0001.

To investigate the marmoset-specific NogoA+ astrocytes further, we extended our analysis to several key pathophysiological time points post-stroke. We were able to demonstrate a transient upregulation of NogoA on GFAP+ reactive astrocytes for at least two weeks post-ischemic stroke in the marmoset (**Supplementary Fig. 4**). We also demonstrated that the population of NogoA+ reactive astrocytes upregulated select markers, such as GAP43 (**Fig. 2b-c**), CD44 and KLF6 (**Supplementary Fig. 3**), in tissue, consistent with our transcriptomic data.

To determine if the NogoA+ reactive astrocytes observed in marmosets were obtained to human stroke, post-mortem human cortical tissue (one-week post-ischemic stroke) was sourced from the Newcastle Brain Tissue Resource (UK). Immunolabeling revealed a subpopulation of GFAP+ reactive astrocytes co-expressing NogoA adjacent to the ischemic core (**Fig. 3e**), consistent with our data in the marmoset at the same pathophysiological time point. Importantly, the expression profile of NogoA in human reactive astrocytes is more consistent with the marmoset and unlikely to be accumulation of myelin debris. These results confirm that the expression of NogoA was associated with astrocytes at an identical time-point post-ischemic stroke in both marmoset and human.

Following CNS injury, myelin breakdown occurs and results in the accumulation of myelin debris proximal to the injury site^17^. Astrocytes are capable of phagocytosis of synapses, debris and dead cells in the CNS^4^. We further determined that NogoA labeling on GFAP+ reactive astrocytes was due to expression and not a consequence of intracellular myelin debris by culturing mouse, marmoset and human astrocytes in the absence of myelin. Mouse, marmoset and human astrocytes expressed NogoA *in vitro* (**Supplementary Fig. 5a**). Astrocytes treated with a pro-inflammatory cytokine, IL-6, exhibited upregulated expression of NogoA (**Supplementary Fig. 5b**). Therefore, these findings confirm that NogoA is upregulated on GFAP+ reactive astrocytes as a direct consequence of stroke and not as a consequence of myelin debris uptake.

### Infiltrating peripheral macrophages in the ischemic zone express NogoA immune-receptor LILRB2

Following ischemic stroke, the blood-brain barrier (BBB) breaks down, permitting infiltration of blood-borne (peripheral) macrophages into the parenchyma^18^. In parallel, reactive astrocytes can exert inhibitory effects on infiltrating inflammatory cell populations in order to mitigate damage following CNS injury, such as producing anti-inflammatory cytokines and corralling immune infiltrates in a specific region^19, 20^. Following ischemic stroke, infiltration of peripheral macrophages peaks around 5-7 DPI in primates, including humans^21, 22^ (**Supplementary Fig. 6a**). Consistent with this, we observed that almost all macrophages at 7 DPI were of peripheral origin, as indicated by the abundant expression of the macrophage marker Iba1 and absence of the microglia-specific marker TMEM119^23^ (**Fig. 4a, c**), which is in line with previous work in the marmoset^2^. Double labeling confirmed the presence of TMEM119-/ Iba1+ peripheral macrophages within the ischemic zone at 7 DPI (**Fig. 4b**). By 21 DPI, we observed the overall absence of peripheral macrophages within the ischemic zone (**Fig. 4c**). Our results indicate that peripheral macrophages infiltrate the ischemic zone parenchyma by 7 DPI, coinciding with the expression of NogoA on reactive astrocytes.

**Figure 4.**
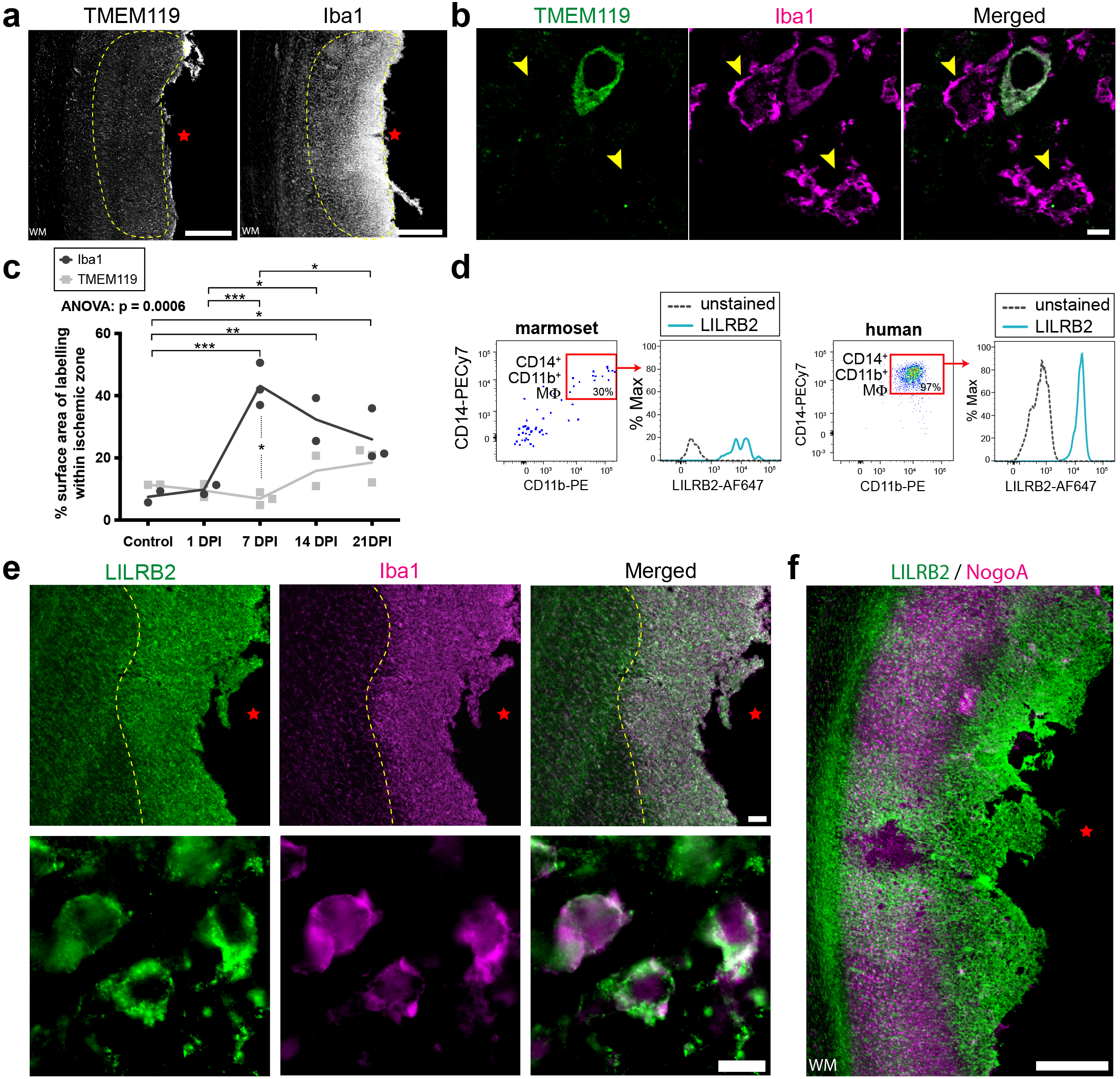
Peripheral macrophage infiltration in the marmoset post-ischemic stroke. (**a**) Representative low-power photomicrographs revealing the low TMEM119 (microglia-specific) and high Iba1 (macrophages) expression within the ischemic zone of marmoset one-week post-ischemic stroke cortical tissue by DAB immunohistochemistry. (**b**) High-power photomicrographs identify a lack of TMEM119 colocalization with Iba1+ cells in the ischemic zone of one-week post-ischemic stroke marmoset. (**c**) Quantification of peripheral macrophage infiltration in the marmoset post-ischemic stroke. Each dot represents the mean surface area labelled with Iba1 versus TMEM119, calculated from a minimum of 3 sections per marmoset, across various post-ischemic stroke recovery time points. (**d**) Marmoset and human macrophage analysis. Fluorescent-activated cell sorting data depicting NogoA immune-receptor (LILRB2) expression in marmoset and human CD14+/ CD11b+ macrophage populations. (**e**) Representative low and high-power photomicrographs demonstrating LILRB2 colocalization with Iba1+ cells in the ischemic zone of one-week post-ischemic stroke marmoset. (**f**) Representative pseudo colored LILRB2 and NogoA photomicrographs overlaid to show spatial distribution of labelling. TMEM119: transmembrane protein 119; Iba1: ionized calcium-binding adapter molecule 1; WM: white matter; yellow dotted line: cell population of interest with lack of TMEM119 expression; red star: ischemic core; yellow arrowheads: TMEM119-/ Iba1+ peripheral macrophages. DPI: days post-ischemic stroke; MØ: macrophages; LILRB2: leukocyte immunoglobulin-like receptor B2; NogoA: neurite outgrowth inhibitor A; scale bars: 500μm (**a, f**), 5μm (**b**), 100μm (**e**: top), 10μm (**e**: bottom); statistical tests: ordinary two way-ANOVA with Sidak’s multiple comparisons; *: p<0.05; **: p<0.01; ***: p<0.001.

Outside the CNS, the NogoA immune-receptor LILRB2 is primarily expressed on mononuclear phagocytes, B cells and dendritic cells^24^. Blood-borne macrophages lacking LILRB2 are hyper-adhesive and spread more rapidly, indicating that LILRB2 functionally limits adhesion and cell spreading^25^. While expression of LILRB2 is known on human peripheral monocytes^24^, the expression on marmoset monocytes is unknown. Based on these data, we asked whether infiltrating macrophages expressed LILRB2 in marmosets and whether macrophage distribution relative to NogoA+ reactive astrocytes was consistent with a receptor-ligand interaction between cell populations. To address these questions, blood was collected from adult marmosets and mononuclear cells were isolated and cultured. Marmoset macrophages were identified as CD14+/ CD11b+ cells^26^, and expression of LILRB2 was detected after 7 days in culture, consistent with humans (**Fig. 4d**). Thus, blood-borne macrophages highly express LILRB2 in the marmoset, as in humans, at baseline.

To determine whether the aforementioned infiltrating peripheral macrophages were expressing LILRB2, we immunolabeled tissues and found a dense distribution of LILRB2+/ Iba1+ macrophages in the ischemic zone at 7 DPI (**Fig. 4e**), compared to controls and other post-ischemic stroke time points (**Supplementary Fig. 6b**). The population of LILRB2+/ Iba1+ cells exhibited a spherical/ amoeboid morphology at 7 DPI, indicative of infiltrating peripheral macrophages^27^ or phagocytic microglia^28^ (**Fig. 4e**). The LILRB2+/ Iba1+ macrophages (**Fig. 4e, yellow-dotted line**) and TMEM119-/ Iba1+ (infiltrating peripheral) macrophages (**Fig. 4a, b, yellow-dotted line**) were localized to the same region of the ischemic zone, indicating that they were the same population of cells. Furthermore, at 7 DPI we observed that deeper infiltrating Iba1+ macrophages had reduced LILRB2 expression and exhibited ramified morphologies compared to the macrophages more proximal to the cortical surface, which highly expressed LILRB2 and were spherical/ amoeboid in morphology (**Supplementary Fig. 6c**). At 7 DPI, the spatial distribution of LILRB2 and NogoA expression were juxtaposed at the ischemic zone (**Fig. 4f**). These results demonstrate that LILRB2 is upregulated on a population of Iba1+ peripheral macrophages 7 DPI with complementary expression to NogoA upregulation on GFAP+ astrocytes. This juxtaposition suggests a potential ligand-receptor interaction contributing to astrocytic corralling of macrophages.

### NogoA-positive reactive astrocytes induce macrophage repulsion through two functional domains

Genetic deletion of the mouse homologue of LILRB2, PirB, after stroke results in an attenuated reactive astrocyte response^29^, indicating a molecular role for LILRB2 related to astrocytes after injury. Additionally, NogoA signaling has been previously shown to limit the migration of microglia *in vitro*^30^. We considered NogoA / LILRB2-dependent immune regulation following CNS injury. The function of NogoA / LILRB2 signaling on peripheral macrophages was investigated using a human monocyte cell line, THP-1-derived macrophages. Analysis of previous transcriptomic data from Gosselin, et al.^23^ revealed the enrichment of LILRB2 on human monocytes (**Fig. 5a**), with an absence of other NogoA binding partners, such as NgR1 and S1PR2. We confirmed this in THP-1-derived macrophages (**Fig. 5b**). LILRB2+ THP-1-derived macrophages, co-expressing Iba1, were phagocytic and exhibited expected morphologies (**Fig. 5c**), including but not limited to multi-nucleation, granulation and heterogeneity in cell shape and size, as described^31^. We subsequently investigated if the two potent functional domains of NogoA -Nogo-66 and Nogo-Δ20 (**Fig. 5d**), could activate LILRB2 signaling on THP-1-derived macrophages by examining key downstream effectors: POSH, Shroom3, RhoA and ROCK1. Nogo-66 treatment induced significant elevation of LILRB2 and POSH at 120 min (**Fig. 5e, g**). Shroom3 and RhoA elevation were detected more acutely at 60 and 30 min, respectively, returning to control levels at subsequent time points analyzed (**Fig. 5e, g**). Nogo-Δ20 treatment induced significant elevation of LILRB2 at 60 min, which was sustained up to 120 min (**Fig. 6f, h**). POSH elevation was detected more acutely starting at 10 min and remaining elevated (**Fig. 5f, h**). Shroom3 and RhoA were elevated at 120 min (**Fig. 6f, h**). No significant changes in ROCK1 expression levels were observed up to 120 min of stimulation with Nogo-66 or Nogo-Δ20 (**Fig. 5e-h**). These data suggest that NogoA induces LILRB2 signaling through the POSH/ Shroom3-dependent pathway in human macrophages via its two distinct functional domains either directly or indirectly. Finally, stripe assays were performed to demonstrate a functional role for NogoA/ LILRB2 signaling in THP-1 human macrophages. Stimulation with either Nogo-66 or Nogo-Δ20 revealed a significantly greater number of Iba1+ macrophages between immobilized human NogoA stripes compared to immobilized control-Fc stripes (**Fig. 5i, j**). Our results demonstrate that LILRB2 blockade significantly attenuates the effect of NogoA signaling on human macrophages resulting in a macrophage distribution indistinguishable from controls (**Fig. 5i, j**). These data demonstrate for the first time that NogoA/ LILRB2 signaling leads to cell-repulsion. Taken together, these data provide substantive evidence that NogoA expression on GFAP+ reactive astrocytes contributes to the corralling of LILRB2+ infiltrating monocytes and macrophages in the primate brain post-ischemic stroke.

**Figure 5.**
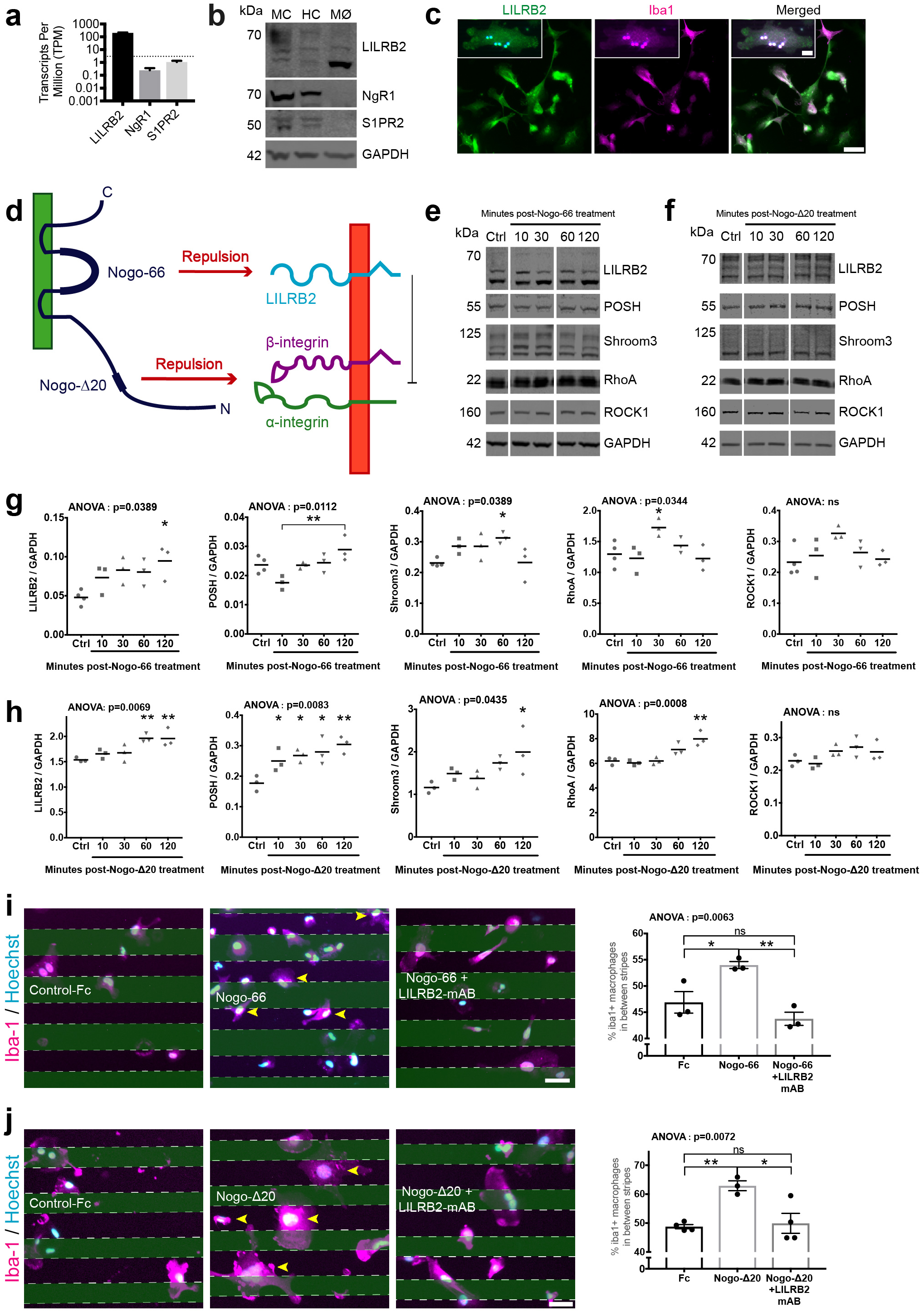
NogoA/ LILRB2 signaling induces repulsion of human macrophages. (**a**) Human RNA seq data from Gosselin *et al.* ^23^ demonstrating known NogoA ligand binding partners in transcripts per million detected in human monocytes. (**b**) Representative immunoblots for LILRB2, NgR1 and S1PR2 in marmoset (MC) and human (HC) cortical tissues, and THP-1-derived human macrophages (MØ). (**c**) Representative LILRB2/ Iba1 immunostaining of THP-1-derived human macrophages, *in vitro*. THP-1-derived human macrophages were tested for phagocytosis of latex beads (**c**, magnified box). (**d**) Schematic depicting NogoA domains and receptors. (**e, f**) Representative immunoblots for THP-1-derived human macrophages treated with human Nogo-66 (**e**) or Nogo-Δ20 (**f**) for 0, 10, 30, 60 and 120 minutes. (**g, h**) Scatter plots depict densitometric quantification of protein normalized to GAPDH. Each dot represents the mean of 3 technical replicates for independent THP-1-derived macrophage cohorts analyzed. (**i, j**) Representative immunostaining of the in vitro stripe (repulsion) assay for human Control-Fc, NogoA and NogoA with LILRB2 blocker pre-treatment of macrophages. The scatter plots with bars depict mean counts with SEM for Iba1+ cells situated in between stripes, n=3. LILRB2: leukocyte immunoglobulin-like receptor B2; RTN4R/ NgR1: Nogo Receptor 1; S1PR2: sphingosine-1-phosphate receptor 2; GAPDH: glyceraldehyde 3-phosphate dehydrogenase; kDa: kilodalton; MØ: macrophages; Iba1: ionized calcium-binding adapter molecule 1; POSH: plenty of sarcoma homology domain 3; ROCK1: Rho kinase 1; NogoA: neurite outgrowth inhibitor A; mAB: monoclonal antibody; yellow arrowheads: NogoA-repelled cells; scale bars: 50μm (**c, i, j**), 10μm (**c**, magnified box); statistical tests: ordinary one way-ANOVA with post-hoc Dunnett’s OR Tukey’s multiple comparisons; *: p<0.05; **: p<0.01; ns: not significant.

Although Nogo-Δ20 is not able to pull down LILRB2 from human macrophage cell lysates (**Supplementary Fig. 7a**), blocking LILRB2 overcomes Nogo-Δ20-mediated repulsion *in vitro* (**Fig. 5j**). Nogo-Δ20 has not been shown to signal through LILRB2 directly, but rather various integrins^32^. As a negative regulator of integrins, the absence or blockade of PirB in macrophages can cause enhanced integrin signaling, resulting in hyper-adhesive macrophages^25^. Therefore, we hypothesize that blocking LILRB2 in our experimental paradigm activated integrin-mediated signaling which overcame Nogo-Δ20-mediated repulsion. This hypothesis is supported by evidence demonstrating that activating integrin-β1 could overcome Nogo-Δ20-mediated repulsion^32^, and our own experiments demonstrating that integrin-α4 and -β3 are significantly upregulated on human macrophages following treatment with a LILRB2 blocking antibody (**Supplementary Fig. 7b**). Thus, these data demonstrate that reactive astrocyte-mediated repulsion of infiltrating macrophages between 7-14 DPI is mediated through NogoA-LILRB2 signaling in the primate (**Supplementary Fig. 8**).

## Discussion

Here we provide a detailed investigation of the cellular and molecular responses of the NHP brain to ischemic stroke, at single-cell resolution. Deep molecular profiling revealed a remarkable diversity of astrocytes in the NHP brain and dramatic shifts in astrocyte subclusters post-ischemic stroke. Notably, in the post-stroke hemisphere, we did not observe presumptive neuroprotective or neurotoxic subpopulations of astrocytes matching those previously described in rodent models. Rather, we identified for the first time, that RTN4A/ NogoA, a membrane protein previously described as an oligodendrocyte-associated neurite outgrowth inhibitor, is also expressed by mammalian astrocytes and its expression is transiently upregulated by astrocytes adjacent to the ischemic core, including in the human cortex. Finally, we demonstrated that NogoA functions in these reactive astrocytes to corral peripheral macrophages from spreading into healthy adjacent CNS tissue. Our data highlights species-specific differences in astrocyte identify across model organisms and emphasizes the need for models that better recapitulate primate, including human, phenomena for the development of therapeutic treatments.

The most widely accepted role for reactive astrocytes has been to create a glial scar, as a structural and molecular barrier to repair and regeneration^5, 33^. However, astrocyte reactivity is not synonymous with repair inhibition, since reactive astrocytes provide crucial neuroprotection after CNS injuries, e.g.: trophic support^34-36^, attenuate excitotoxicity^13, 37-39^, eliminate cell debris^40^ and, regulate neuroinflammation^19, 41^. Indeed, astrocyte functional diversity may be necessary for axonal regeneration^42^. We observe heterogenous astrocyte identities in the intact contralateral NHP V1, defined by unique gene expression profiles predominantly associated with the regulation of synaptic transmission and myelination. This large diversity is unsurprising considering recent evidence demonstrating regional^43, 44^ and laminae^45^ specific astrocyte identities in the neocortex, highlighting that astrocyte diversity is extensive. This heterogeneity is further enhanced after injury by the emergence of distinct subsets of reactive astrocytes upregulating genes associated with immunomodulation. We demonstrate that NHP reactive astrocytes are not aligned with either the A1 or A2 astrocyte profiles previously identified in rodents^5, 6^. Indeed, while a minor subset of A1 markers (MX1, SerpinG1, and FKBP5) were enriched in several reactive astrocyte subtypes, most A1 markers analysed were not, including crucial indicators such as PSMB8, SRGN and especially complement component C3, previously described as a surrogate marker of A1 astrocytes^5, 46-48^. Instead, all reactive astrocyte subtypes upregulated a combination of A1 and A2 markers. While we acknowledge the caveat surrounding the model species/ pathologies necessary to induce these discrete reactive astrocyte subtypes, our data reinforces the hypothesis that astrocyte reactivity undergoes multiple stages and paths of activation. Cataloguing the diversity and plasticity of reactive astrocytes is crucial to fully elucidate the extent to which reactive astrocyte contributes to the process of repair and regeneration after CNS injuries.

Recognizably, reactive astrocyte characteristics significantly differ between rodents and humans, including diverging microanatomical^9^ and transcriptional characteristics^49^, which collectively underpin functional differences. The failure to observe NogoA on reactive astrocytes over the past decades is likely due to these interspecies gene expression differences in astrocytes. A recent study directly comparing primary resting astrocytes derived from mouse versus human brain, demonstrated that that more than 600 genes enriched in human astrocytes were not similarly enriched in mouse astrocytes^49^. Furthermore, extrinsic factors that influence astrocyte gene expression and subsequent function, such as the immune response, differ significantly between between rodents and primates. For example, human and mouse macrophages respond very differently to the same stimuli^50^, where in response to lipopolysaccharide, mouse macrophages upregulate INOS, human macrophages characteristically upregulate CCL20, CXCL13, IL-7R, P2RX7, and STAT4^51^. The combination of these interspecies differences is likely to culminate in variations of/ additional mechanisms in the interplay between astrocytes and monocytes/ macrophages after CNS injury. As such, the enrichment of NogoA on a large proportion of reactive astrocytes in primates, including human, but not rodents, which coincides temporally with peripheral macrophage infiltration after ischemic stroke, is likely a consequence of requiring a more complex reactive astrocyte response to a more complex immune response. The functional addition of NogoA to astrocyte-mediated corralling of infiltrating leukocytes reinforces the physiological importance of this neuroprotective mechanism to spare healthy brain tissue and confine the injury to a discrete region.

With regard to therapeutic intervention, NogoA and other myelin-associated inhibitors have been extensively studied in their capacity as inhibitors of repair and have been identified as promising targets for therapeutic intervention, especially for the treatment of SCI^52-61^ and stroke^62-68^. A NogoA neutralising antibody, IN-1, has been tested in rodent models of SCI^52, 54, 56, 69^, followed by NHPs^58, 70, 71^, and finally human subjects^72^. The Phase I clinical trial using ATI335 (NogoA neutralizing antibody) in acute paraplegic and tetraplegic patients reported that it is well tolerated following intrathecal administration^72^ and has entered Phase II trials in subacute SCI (ClinicalTrials.gov NCT03935321). In addition, a soluble NgR decoy blocking Nogo, MAG and OMgp ligands is efficacious preclinically in chronic SCI^73, 74^, and has entered clinical trials for that indication (ClinicalTrials.gov NCT03989440).

Importantly, our findings from acute to subacute time points following ischemic stroke in the marmoset indicate that NogoA antagonism might restrict astrocyte-mediated corralling of infiltrating macrophages. In addition, NogoA blockade might have indirect immunomodulatory effects at early time points. Here, we find that the most enriched functional gene categories in NogoA+ astrocytes include wound healing, blood vessel development, regulation of cell adhesion, ECM organization and cytokine-mediate signaling pathway, all of which are immunomodulatory functions. We know that reactive astrocytes physically restrict migration of leukocytes, including macrophages, after CNS injury to limit their infiltration into adjacent healthy tissue^19, 75^. While immune infiltrates are crucial for the clearance of detrimental cell and myelin debris^76^, the inflammatory consequences may exacerbate secondary neuronal cell death. For example, if proliferating or corralling reactive astrocytes are ablated, the result is propagation of leukocyte infiltration, severe demyelination, neuronal and oligodendrocyte cell death and pronounced motor deficits ensue following CNS trauma^77, 78^. Reactive astrocyte expression of the receptors ERα, GP130, DRD2 and C5aR1^79-83^ provides potential alternative mechanisms to limit immune infiltration. However, DRD2 and C5aR1 are absent or negligible in marmoset reactive astrocytes. Only two of these aforementioned astrocyte receptors (ERα & GP130) are significantly upregulated in marmoset reactive astrocytes one-week after stroke, with up to 40% of NogoA+ reactive astrocytes expressing GP130. In primates, NogoA signaling may be crucial to fully restrict macrophage infiltration into surrounding healthy CNS parenchyma after stroke. Therefore, different mechanisms may mediate macrophage corralling in rodents and primates after ischemic stroke. NogoA-directed therapies may titrate both early modulation of inflammation, as well as longer term regulation of axonal sprouting and neural repair.

In sum, our results support that a NHP ischemic stroke model is more representative of human sequelae than the rodent, highlighted by our verification of NogoA+ reactive astrocytes both in marmoset and human cortical tissue one-week post-stroke. Given the complex and species-specific response of astroglial cells to CNS injury, it is essential to complete a detailed molecular characterization of NHP preclinical models, considering the immune system implication amongst others, to advance ischemic stroke research and therapeutic development.

**Online Content**

## Methods

### Animals

Adult common marmosets (*Callithrix jacchus* >18months; n=15 fixed, n=16 fresh, n=3 for transcriptomics) were used in this study. Animals were subdivided into uninjured control and injured cohorts, comprising 1, 7, 14 and 21 days post injury (DPI) recovery periods (fixed: n=3 per time point; fresh: n=3 per time point, excluding 14 DPI, where n=4). n=3 for transcriptomics were allocated to the 7 DPI recovery period with the contralateral hemisphere used as control. This route was chosen to reduce the number of animals sacrificed for our studies. While we acknowledge that there are likely to be distal changes to brain regions following stroke, our focus in this study were the local changes at the site of the stroke given the highly focal nature of our stroke model^2^. Moreover, our in-tissue validation of NogoA, GAP43, CD44 and KLF6 labelling using true control tissues was consistent with our transcriptomic analysis using contralateral tissues. Experiments were conducted according to the Australian Code of Practice for the Care and Use of Animals for Scientific Purposes and were approved by the Monash University Animal Ethics Committee. Animals were obtained and housed at the National Nonhuman Primate Breeding and Research Facility (Monash University).

### ET-1-induced focal ischemic stroke

Preoperative procedures, anesthesia and surgery were all performed on adult marmoset monkeys, as previously described^2^. In short, anesthesia was induced using Alfaxalone (5mg/kg) and maintained using inspired isoflurane (0.5-4%) throughout all surgical procedures. Induction of focal ischemic injury to adult marmoset primary visual cortex (V1) was achieved by vasoconstrictor-mediated vascular occlusion of the calcarine branch of the posterior cerebral artery (PCAca), which supplies operculum V1. Following midline incision, craniotomy and dural resection, intracortical injections of endothelin-1 (ET-1: 1mg/mL; rate: 0.1µL/30s pulse + 30s intervals, totaling 0.7µL over 7 sites) proximal to the PCAca were performed. Upon completion of injections, the craniotomy was replaced and positioned with tissue adhesive (Vetbond 3M) and the skin sutured closed. Uninjured animals were used as controls. Stroke model is summarized in **Fig. 1a** and **Supplementary Fig. 1a**.

### Single nuclei 10x genomic sequencing and analysis

#### Sample collection

One-week post-ET-1-induced ischemic stroke in V1, adult marmosets (n=3) were administered an overdose of sodium pentobarbitone (100mg/ kg; IM). Following the loss of corneal and muscular reflexes, the animals were decapitated, and the brain removed under aseptic conditions. Cerebral tissues were rinsed in chilled PBS, to remove excess blood, before dissection of regions of interest with the aid of a dissecting microscope (**Fig. 1a**; Created with BioRender.com). Dissected tissues were placed in sterile tubes and dropped into liquid nitrogen before storage at −80°C. The procedures/ dissections were performed in chilled RNAase-free PBS with RNase-free sterilized instruments under RNase-free conditions. Approximate time from apnea to snap-freezing ranged from 20-30mins. 4 samples per animal = 12 samples for analysis. All 12 samples passed QC.

#### Brain Cell Nuclei Isolation

Frozen marmoset brain region tissue was finely pulverized to powder in liquid nitrogen with mortar and pestle (Coorstek #60316, #60317). All buffers were ice-cold and all reagents used for consequent nuclear isolation were molecular biology grade unless stated otherwise. 50 mg of pulverized tissue was added into 5 mL of ice-cold lysis buffer: 320 mM sucrose (Sigma #S0389), 5 mM CaCl_2_ (Sigma #21115), 3 mM Mg(Ace)_2_ (Sigma #63052), 10mM Tris-HCl (pH 8) (AmericanBio #AB14043), protease inhibitors w/o EDTA (Roche #11836170001), 0.1 mM EDTA (AmericanBio #AB00502), RNAse inhibitor (80U/mL) (Roche #03335402001), 1mM DTT (Sigma #43186), 0.1% TX-100 (v/v) (Sigma #T8787). Reagents: DTT, RNAse Protector, protease inhibitors, TX-100 were added immediately before use. The suspension was transferred to Dounce tissue grinder (15mL volume, Wheaton #357544; autoclaved, RNAse free, ice-cold) and homogenized with loose and tight pestles, 30 cycles each, with constant pressure and without introduction of air. The homogenate was strained through 40 um tube top cell strainer (Corning #352340) which was pre-wetted with 1mL isolation buffer: 1800 mM sucrose (Sigma #S0389), 3 mM Mg(Ace)_2_ (Sigma #63052), 10mM Tris-HCl (pH 8) (AmericanBio #AB14043), protease inhibitors w/o EDTA (Roche #11836170001), RNAse inhibitor (80U/mL) (Roche #03335402001), 1mM DTT (Sigma #43186). Additional 9 mL of isolation buffer was added to wash the strainer. Final 15 mL of solution was mixed by inverting the tube 10x and carefully pipetted into 2 ultracentrifuge tubes (Beckman Coulter #344059) onto the isolation buffer cushion (5 mL) without disrupting the phases. The tubes were centrifuged at 30000 x *g*, for 60 min at 4 °C on ultracentrifuge (Beckman L7-65) and rotor (Beckman SW41-Ti). Upon end of ultracentrifugation, the supernatant was carefully and completely removed and 100 ul of resuspension buffer (250 mM sucrose (Sigma #S0389), 25 mM KCl (Sigma #60142), 5mM MgCl2 (Sigma #M1028), 20mM Tris-HCl (pH 7.5) (AmericanBio #AB14043; Sigma #T2413), protease inhibitors w/o EDTA (Roche #11836170001), RNAse inhibitor (80U/mL) (Roche #03335402001), 1mM DTT (Sigma #43186)) was added dropwise on the pellet in each tube and incubated on ice for 15 minutes. Pellets were gently dissolved by pipetting 30x with 1mL pipette tip, pooled and filtered through 40 um tube top cell strainer (Corning #352340). Finally, nuclei were counted on hemocytometer and diluted to 1 million/mL with sample-run buffer: 0.1% BSA (Gemini Bio-Products #700-106P), RNAse inhibitor (80U/mL) (Roche #03335402001), 1mM DTT (Sigma #43186) in DPBS (Gibco #14190).

#### Single cell and/ or nucleus microfluidic capture and cDNA synthesis

The cells and nuclei samples were placed on ice and taken to Yale Center for Genome Analysis core facility and processed within 15 minutes for single nucleus RNA sequencing with targeted nuclei recovery of 10000 nuclei, with Chromium Single Cell 3’ v3 Chemistry (10x Genomics) on microfluidic Chromium System (10x Genomics) by following precisely manufacturers detailed protocol (CG000183_ChromiumSingleCell3’_v3_UG_RevA). Specifically, due to limitations imposed by source RNA quantity, cDNA from nuclei was amplified for 14 cycles.

#### Single cell and/ or nucleus RNAseq library preparation

The post cDNA amplification cleanup and construction of sample-indexed libraries and their amplification subsequently precisely followed manufacturer’s directions (CG000183_ChromiumSingleCell3’_v3_UG_RevA). Specifically, the amplification step directly depended on the quantity of input cDNA and it varied from 8 – 14 cycles.

#### Sequencing of libraries

In order to reach optimal sequencing depth (25,000 raw reads per nucleus), single cell and/or nucleus libraries were run using paired-end sequencing with single indexing on the HiSeq 4000 platform (Illumina) by following manufacturer’s instructions (Illumina, 10x Genomics) (CG000183_ChromiumSingleCell3’_v3_UG_RevA). To avoid lane bias, multiple uniquely indexed samples were mixed and distributed over several lanes.

#### Cell de-multiplexing, reads alignment and gene expression quantification

The *cellranger*^84^ commercial software was utilized to conduct the initial data processes. First, the *cellranger mkfastq* was used to demultiplex the raw base call (BCL) files obtained from Illumina HiSeq 4000 sequencers into FASTQ files. Second, the *cellranger mkref* was used to build a reference index for the marmoset reference genome (calJac3), and the GTF format Ensembl^85^ gene annotation (Callithrix_jacchus.C_jacchus3.2.1.91.gtf) was processed to the transcriptional model specified for single nuclei sequencing. Third, the *cellranger count* was used to perform read alignments to the reference genome, cell filtering and counting, and gene UMI quantification for every single cell. Notably, all parameters of *cellranger* were set as default, except for “--expect-cells = 10000”.

#### Removal of nuclei doublets

Nuclei doublets or clumps were removed through post-transcriptomic analysis, using *Scrublet86* software with a filtered feature barcode matrix (i.e., *matrix.mtx*) generated by *cellranger* as input. To optimize the analysis, we customized some parameters based on in-house computing trials, i.e., *expected_doublet_rate=0.05, min_counts=2, min_cells=3, min_gene_variability_pctl=85, n_prin_comps=30.* The removal of nuclei doublets was conducted separately for each sample, from which cross-sample variation was observed, although the rates of nuclei doublets were all less than 11% (**Supplementary Fig 1c**).

#### Data analysis and mining

*Seurat*^87^ and some in-house customized scripts were used to manage the single nuclei transcriptomic dataset. Specifically, raw gene UMI counts per nuclei were used to create a *Seurat* object followed by log normalization using *NormalizeData* function with setting parameters *normalization.method = “LogNormalize”, scale.factor = 10000*. To improve the signal-to-noise ratio, and produce robust and reliable results, we first identified highly variable genes (HVGs) using *Seurat FindVariableFeatures* function by choosing parameters *selection.method = “vst”, nfeatures=3000*. To achieve this goal, we fit the relationship between log-transformed variance and log-transformed mean of gene UMI counts using local polynomial regression, and then chose the 3000 top ranked genes as HVGs, which were scaled subsequently to feed the request from gene dimension reduction and cell clustering. For gene dimension reduction, the first step used principle component analysis (PCA) via a linear dimension reduction approach, which was implemented by using the *Seurat RunPCA* function and consequently choosing the top 50 principal components (PCs) to represent cell variation. The second step used uniform manifold approximation and projection (UMAP)^88^ to complete further dimension reduction of the top 30 PCs generated in the first step. Finally, we used the *Seurat FindClusters* function with parameters (i.e., *dims=1:30, resolution=1*) to analyze nuclei clustering. Notably, these analyses were repeated at least twice. The first round aimed at identifying major nuclei clustering that would ultimately be associated with the main cell type. The second round took the main cell types and divided nuclei into subtypes with higher resolution.

#### Assignment of cell type

In order to associate nuclei clusters to specific cell types, we calculated the gene specificity score for each gene in each cluster using the R script retrieved from Efroni, et al. ^89^. Genes with the highest specificity score suggested enrichment and were consequently considered as cluster-specific markers, which would be compared with the literature of reported cell type markers to assign a cell type to each cluster. Cell-type markers were as follows: astrocyte (SLC1A2, SLC1A3, ALDH1L1, GFAP, AQP4, NDRG2), endothelial cell (FLT1, IGFBP7, SLC2A1), excitatory neuron (SYT1, RBFOX3, GRIN2A, SATB2, CUX1, CUX2), inhibitory neuron (GAD1, GAD2, LHX6, SST, VIP), microglia (P2RY12, CD74, DOCK8, GPR34, C1QB), oligodendrocyte (PLP1, MBP, UGT8, ST18, MOBP), and oligodendrocyte precursor cell (PDGFRA, PCDH15, MEGF11). Nuclei clusters enriched with a particular set of markers were considered to be of the corresponding cell type.

#### Removal of unwanted batch effects

In order to reduce unwanted batch effects that originated from external systematic or technical issues, we utilized Fast Mutual Nearest Neighbors (*fastMNN*)^90^ correction. Here, we considered the three different marmosets as the potential source of external batches, since the sample dissection and library preparation were separately executed; there was also the possibility that individual differences could be a contributor. Instead of using fastMNN directly, we used the *RunFastMNN* function, which is efficiently integrated in the *SeuratWrappers*^91^ package, to unite this analysis to the whole pipeline. Additionally, the execution of the *RunFastMNN* function was based on some HVGs, which were identified separately but shared by the three batches, using *Seurat FindVariableFeatures* function with parameters *selection.method = “vst”, nfeatures=3000*. Notably, the removal of unwanted batch effects was conducted separately for each brain region and each cell type.

#### Differential expression analysis and gene ontology analysis

The *Seurat FindAllMarkers* function was adopted to identify differentially expressed genes. In brief, we took one group of nuclei and compared it with another group of nuclei, using a *Wilcox* model. In addition, the parameters “*logfc.threshold*” and “*min.pct*” were set to 1e-5 to strengthen the power for comparing genes with less signal. For detecting top candidates, we defined statistically significant as genes that had greater than 0.25 log(e)-transformed fold change between groups in either direction and exhibited adjusted p-values (False Discovery Rate) less than 0.01; further, only genes that were expressed by at least 10% of nuclei in either population were considered. Specific to the analysis of injured versus contralateral nuclei, we also considered genes to be statistically significant if the ratio of nuclei expressing the gene between inj and con nuclei was >1.25, even if the fold change was below our threshold.

Differentially expressed gene lists were analyzed using the Express Analysis function of Metascape^92^. To assess functional enrichment in the individual astrocyte subtypes, and between the injured and contralateral conditions, the top 50 genes were used (based on specificity score or FDR-adjusted p-value, respectively). To identify enriched processes between NogoA-positive and NogoA-negative astrocytes, all DEGs (131 genes) were used.

### Fresh and fixed tissue collection for protein analyses

At the end of the designated post-stroke period: 1 DPI, 7 DPI, 14 DPI or 21 DPI; marmosets were administered an overdose of pentobarbitone (100mg/kg; IM). For fresh tissue, brains were immediately harvested following apnea. The occipital poles were dissected at the level of the diencephalon and bisected coronally, with caudal portions encompassing V1 and the ischemic zone and core, before snap freezing them in liquid nitrogen. For fixed tissue, animals were similarly euthanized and transcardially perfused with 0.1M heparinized saline solution (0.9% sodium chloride at 37°C containing 0.1M heparin), followed by 4% paraformaldehyde. Brains were collected, post-fixed for 24h in 4% paraformaldehyde, sucrose protected and cryosectioned, as previously outlined^2, 93^.

### Antibody characterization

NogoA and LILRB2 antibodies (**Supplementary Table 4**) were tested in rat, marmoset and human cortical tissue to ensure uniformity of detectable bands across species (**Supplementary Fig. 9a**). NogoA antibody specificity was validated using two antibodies recognizing separate epitopes on the NogoA ligand due to the ambiguity of some of the data surrounding NogoA expression in various CNS injury models. The NogoA antibodies were tested on fresh and fixed marmoset cortical tissue, rat cortical tissue as a positive control and marmoset liver as a negative control, with and without pre-incubation with blocking peptides to determine antibody specificity (**Supplementary Fig. 9b, c**). TMEM119 and LILRB2 antibodies (**Supplementary Table 4**) were tested in fixed marmoset spleen and brain tissue with an Iba1 co-label to ensure specificity of TMEM119 to microglia in marmoset (**Supplementary Fig. 9d**).

### 3 days post-MCAO mouse tissue

C57Bl6/J mouse cortical tissue (n=3) was gifted from Professor Christopher G. Sobey. 8-12 week old mice were subjected to transient MCAO for 1hour and brains were collected 72 hours post-ischemia. Unperfused fresh brains were snap frozen over liquid nitrogen and sectioned onto slides at 10μm thick and 420μm apart before storing at −80°C. Sections were post-fixed in 4% PFA before immunostaining as described in above protocols.

### Human tissue

Snap-frozen and paraffin-fixed human brain samples (n=1; Age: 74yrs; Gender: Female; COD: Cirrhosis; Post-mortem processing: <24hrs) were obtained from the Newcastle Brain Tissue Resource (UK). Procurement and use of human brain tissue were approved by the Monash University Human Research Ethics Committee in compliance with section 5.1.22 of the National Statement on Ethical Conduct in Human Research (Ethics approval number: CF14/2120 - 2014001121).

### Cell culture

#### Astrocytes

Astrocytes were obtained from mouse, marmoset and human. For mouse astrocytes, magnetic isolation of ASCA-2+ astrocytes was performed. ASCA-2+ astrocytes were purified by immunopanning from seven P5 mouse pup cortices. Briefly, cortices were dissociated mechanically (using a scalpel blade) and enzymatically (using Papain: Worthington, LS003126) with various triturations and Nitex (Sefar, 03-20/14) filtration steps to generate a single-cell suspension. The cell suspension was subsequently allowed to recover at 37°C for 30–45 min in a 10% CO_2_ incubator. The cell suspension underwent debris (Miltenyi, 130-109-398) and red blood cell (Miltenyi Cat. 130-094-183) removal before incubation to positively select for astrocytes using anti-ASCA-2 MicroBead Kit with FcR Blocking Reagent (Miltenyi, 130-097-679) and an MS column/ MACS separator. Isolated astrocytes were cultured at 37°C in a 10% CO_2_ tissue-culture incubator in complete Astrocyte Growth Medium (AGM) containing: 50% Neurobasal medium (Life Technologies, 21103049), 50% Dulbecco’s modified eagle medium (DMEM, Life Technologies, 11960044), 100 units of penicillin and 100µg/mL streptomycin (Life Technologies, 15140122), 1 mM Sodium Pyruvate (Life Technologies, 11360070), 292 µg/mL L-glutamine (Life Technologies, 25030081), 5µg/mL N-Acetyl-L-cysteine (NAC, Sigma, A8199), 1x SATO and 5ng/mL HB-EGF (Peprotech, 100-47). The culture was maintained by replacing 50% of the medium every 7 days with complete AGM containing fresh 5ng/mL HB-EGF.

For marmoset astrocytes, postnatal day 14 marmoset V1 was processed as previously described^94^. For human astrocytes, a primary astrocyte cell line was obtained from Lonza. Human astrocytes were maintained in AGM and treated in DMEM serum-free conditions with IL-6 and IL-6 receptor for 24 hours to induce a more reactive phenotype.

#### Macrophages

Human macrophages, the THP-1 monocytic cell line (ATCC® TIB202(tm)), were cultured in RPMI 1640 medium (ATCC® 302001(tm)), supplemented with 0.05mM 2•mercaptoethanol and 10% fetal bovine serum (FBS) in antibiotic-free conditions. THP-1 cells were then differentiated into macrophages with 0.1µM 1α,25-dihydroxyvitamin D3 (vitamin D3; Sigma-Aldrich) or 200nM phorbol 12-myristate 13-acetate (PMA; Sigma-Aldrich) over 72 hours, as previously described^31^. Cells were then treated with 200ng/mL of recombinant rat NogoA (1026-1090aa) and Fc Chimera protein (Nogo-66; R&D Systems), or with human NogoA (566-748aa) and Fc chimera protein (Nogo-Δ20; R&D Systems), for 10m, 30m, 1h and 2h.

### Protein extraction, SDS-PAGE and immunoblot

Snap frozen V1 tissues were homogenized in either TRIzol LS Reagent (ThermoFisher), or NP40 (ThermoFisher) lysis buffer, and protein extracted according to manufacturer’s instructions. Subsequent lysates were supplemented with 10% Protease Inhibitor Cocktail (PrIC; Sigma-Aldrich), 1% Phosphatase Inhibitor Cocktail 2 (PhIC; Sigma-Aldrich), 1mM PMSF Serine Protease Inhibitor (Sigma-Aldrich). Protein concentration was determined using Bradford Reagent (Sigma-Aldrich). At experimental endpoints, cells were lysed using RIPA buffer (ThermoFisher) and addition of previously mentioned inhibitor cocktails. Marmoset and human frontal cortical tissues were also lysed in this way for use as positive controls. Equal concentrations of samples were added to 4X Loading Buffer (240nM TRIS, 8% SDS, 40% glycerol, 20% 2-mecaptoethanol, 0.05% bromophenol blue), heated at 95°C for 10 mins, and electrophoretically separated on a 4-12% Bis-Tris gel (ThermoFisher). Following gel electrophoresis, proteins were blotted onto PVDF membranes (ThermoFisher), pre-blocked in Odyssey Blocking Buffer (PBS; Li-Cor) before incubation with primary antibodies (**Supplementary Table 4**) overnight at 4°C. Following washes, membranes were incubated with IRDYE secondary antibodies (Li-Cor) and visualized using the Odyssey CLx Scanner (Li-Cor).

### Densitometry

Densitometric analysis was performed on scanned immunoblots using Image Studio Lite (Li-Cor: version 5.2.5), and values representing protein over loading control were calculated. Data distribution was evaluated for Gaussian distribution using the Shapiro-Wilk normality test. Ordinary one way-ANOVA with Dunnett’s multiple comparisons post-hoc test were used for statistical analysis of densitometry data - GraphPad Prism 7 (GraphPad). ImageJ ^95^ and Adobe Illustrator CC 2017 (Adobe) were used for post-processing of immunoblot images and figure design.

### Immunohistochemistry and immunofluorescence

For immunohistochemistry (IHC), free-floating sections representing V1 were treated with 0.3% hydrogen peroxide and 50% methanol in 0.01M PBS for 30 minutes to inactivate endogenous peroxidases before pre-blocking. 15% normal horse serum (NHS; Gibco, ThermoFisher, USA) in 0.01M PBS, 0.3% TritonX-100 (PBS-TX; Sigma-Aldrich) was used to pre-block tissue for IHC and immunofluorescence (IF) before incubation with primary antibodies (**Supplementary Table 4**) overnight at 4°C. For secondary labeling sections were incubated with either biotinylated secondary antibodies or Alexa Fluor donkey anti-host secondary antibodies for 1 hour at room temperature. Following secondary incubation, sections for IHC were treated with streptavidin-horseradish peroxidase conjugate (GE Healthcare, Amersham, UK; 1:200) prior to visualization using metal enhanced chromagen, 3,3’-diaminobenzidine (DAB: Sigma-Aldrich). For IF staining, sections were treated with Hoechst 333258 nuclei stain, mounted on Superfrost slides and treated with 0.05% Sudan Black in 70% ethanol for 10mins before coverslipping using fluoromount-G.

For human tissue, sections were dewaxed in xylene before serial rehydration in ethanol. Antigen retrieval was performed using 0.05% citriconic anhydride at 90°C for 2 hours. Sections were subsequently blocked in 5% normal goat serum (NGS; Gibco, ThermoFisher, USA), 1% bovine serum albumin (BSA; Sigma-Aldrich) and 0.1% fish skin gelatin (Sigma-Aldrich) in 0.01M PBS, 2% TritonX-100 (PBS-TX; Sigma-Aldrich) for 1 hour before incubation with primary antibodies: rabbit anti-NogoA (1:200; **Supplementary Table 4**) and mouse anti-GFAP (1:500; Sigma-Aldrich), for 72 hours at 4°C. Secondary labeling was performed using goat anti-rabbit (Alexa Fluor 350; ThermoFisher) and donkey anti-mouse (Alexa Fluor 647; ThermoFisher) overnight at 4°C before coverslipping using fluoromount-G.

For cell culture staining, cells were fixed on poly-ornithine (1mg/mL) and laminin-coated (1mg/mL) glass coverslips or chamber slides in 4% PFA before blocking in 10% NHS in 0.01M PBS, 0.3% PBS-TX. Cells were incubated with primary antibodies (**Supplementary Table 4**) for 2 hours at room temperature before secondary labeling using Alexa Fluor donkey anti-host secondary antibodies for 1 hour at room temperature. Cells were treated with Hoechst 333258 nuclei stain and coverslipped using fluoromount-G.

Stained tissue/cells were analyzed using either the Axioimager Z1 Upright Microscope and Axiovision Software (Carl Zeiss), or the Sp5 Inverted Confocal Microscope, and LAS Software (Leica Microsystems). ImageJ, Adobe Photoshop and Illustrator CC 2017 (Adobe) were used for image post-processing and figure design.

For **Figure 4f** the representative pseudo colored LILRB2 and NogoA photomicrographs were obtained by DAB immunohistochemistry and overlaid using Adobe Photoshop in order to show spatial distribution of labelling. Please note that these photomicrographs are from different (adjacent) sections and have been overlaid to create a representative image as double labelling with LILRB2 and NogoA was not possible at the time due to available antibodies.

### NogoA+/ GFAP+ cell quantification

Z stack images were taken at high magnification of randomly selected GFAP+ cells within 500µm of the ischemic core in 3 days post-MCAO mouse tissue (n=3) and in 7 days post-ischemia marmoset tissue (n=4). Colocalization of NogoA and GFAP or lack thereof was determined using ImageJ and subsequently quantified as the number of NogoA+ astrocytes over the total number of counted astrocytes (GFAP+ cells) and expressed as a percentage. A minimum of 50 cells per animal, over 3-5 sections across 3 mice and 4 marmosets, were imaged and counted. These images were also used to determine the fluorescent intensity of NogoA expression by GFAP+ astrocytes. 10 cells were randomly selected from each of the 3 mice and 4 marmosets, and the mean fluorescent intensity of NogoA was calculated over the mean fluorescent intensity of GFAP using ImageJ and expressed as a ratio of NogoA: GFAP. Data distribution for both cell counts and fluorescent intensity were evaluated for Gaussian distribution using the Shapiro-Wilk normality test. Unpaired t tests were used for statistical analysis of data using GraphPad Prism 7 (GraphPad).

### Iba1/ TMEM119 quantification

Iba1 and TMEM119 DAB-immunolabeled marmoset V1 tissues across control (n=2), 1 (n=2), 7 (n=3), 14 (n=2) and 21 (n=3) DPI time points were imaged and photo-merged using Adobe Photoshop. The area of Iba1 and TMEM119 immunolabeling was quantified over 3 sections from each animal using Image J. Data distribution was evaluated for Gaussian distribution using the Shapiro-Wilk normality test. Ordinary two way-ANOVA and Sidak’s multiple comparisons post-hoc test were used for statistical analysis of data using GraphPad Prism 7 (GraphPad).

### Marmoset and human blood-derived macrophages for flow cytometry

#### Isolation of marmoset blood mononuclear cells

Marmoset blood was collected by cardiac puncture using syringes pre-coated with sodium citrate solution (3.8% w/v) and expelled into citrate coated collection tubes (∼10% v/v of citrate solution: blood volume). Mononuclear cells were collected by density centrifugation using lymphocyte separation media (1.077 g/mL, Lonza) as previously described ^96^. Red blood cells were removed by lysis using NH_4_Cl lysis buffer, the MNC washed twice with PBS (0.5% BSA), filtered through a 40µm strainer and counted.

#### Isolation of human blood mononuclear cells

Human blood was obtained from healthy donors (Australian Red Cross Blood Service, Melbourne) in citrate coated bags. Low density mononuclear cells (MNC) were isolated by discontinuous density centrifugation using Ficoll-Hypaque (1.077 g/mL, Pharmacia Biotech). Red blood cells were removed by lysis using NH_4_Cl lysis buffer, the MNC washed twice with PBS (0.5% BSA), filtered through a 40µm strainer and counted using a CELL-DYN Emerald hematology analyzer (Abbott). All experiments were undertaken following informed consent from donors and approval from the Monash human ethics committee.

#### Autologous marmoset and human serum

To harvest autologous serum, freshly harvested non-fractionated marmoset or human blood was centrifuged at 400 x g and plasma collected. The remaining red/white blood cell layer was processed for MNC the next day as described above. The collected plasma was calcified with 20% (w/v) CaCl_2_ solution in H_2_O at a ratio of 1:100 and 1:20 v/v CaCl_2_ solution: plasma for human and marmoset plasma respectively. The mixture was left undisturbed overnight at 4°C, centrifuged at 2000 x g to remove fibrinogen clots and the serum collected and stored in aliquots at −20°C.

#### In vitro culture of monocyte-derived macrophages

Marmoset or human MNC were resuspended in DMEM (0.5% BSA) at a density of 3 × 10^6^ cells/mL and seeded into 12 or 24-well tissue culture plates. The cells were incubated in a CO_2_ incubator at 37°C for 1 hour prior to non-adherent cells being removed by aspiration, leaving behind adherent monocytes. Fresh DMEM (supplemented with 10% autologous serum and Gibco GlutaMAX) was added and the adherent cells cultured for 5 days (human) or 7 days (marmoset) with autologous serum/media being replaced on day 3 and 5. The plates were gently washed with warm PBS, the adherent macrophages lifted with trypLE Express (Gibco) cell dissociation enzyme and washed with PBS (0.5% BSA) for subsequent analysis.

#### Flow cytometry

For analysis of LILRB2 expression on marmoset and human monocyte derived macrophages were first blocked with human TruStain FcX Fc blocking solution (Biolegend), then labelled with and without anti-human LILRB2 (5 µg/mL; R&D Systems). The cells were washed with PBS (0.5% BSA) and then labelled with goat anti-mouse IgG2a-AF647 (2 µg/mL) secondary antibody. Finally, the cells were washed and labelled with a cocktail containing anti-human CD11b-PE (clone D12; BD Biosciences) and anti-human CD14-PECy7 (clone M5E2; BD Biosciences), washed, resuspended in PBS (0.5% BSA plus propidium iodide) and analyzed on an LSR II (BD Biosciences), as previously described ^97^. Flow cytometric analysis was performed using FlowJo software. The expression of LILRB2 was assessed on macrophages, which were identified as CD14+/ CD11b+ cells that had been pre-gated on single (FSC-H vs FSC-A), nucleated (SSC-A vs FSC-A), viable (PI^neg^) cells.

### Stripe (attraction/ repulsion) assay

13mm diameter glass coverslips were cleaned overnight in 2M HCl at 60°C followed by sonication in graded alcohols. Coverslips were then coated overnight with Poly-L-Ornithine (1mg/mL) at 37°C. Coverslips were dried and placed coated-side down on a silicon matrix with 50µm wide, 50µm apart alternating grooves (obtained from Dr. Martin Bastmeyer, Karlsrube University, Germany). Nogo-66 (10µg/mL), Nogo-Δ20 (10µg/mL), or recombinant human fc (10µg/mL) were injected into the constructs and incubated for 1.5hrs at room temperature in order to create the stripes. PBS was injected to flush and wash out unbound Fc proteins and the coverslips removed from the matrix and placed face-up in a 24-well plate. Coverslips were subsequently washed in PBS followed by laminin (1mg/mL in MEM) for 2 hrs at 37°C. Human THP-1 macrophages (differentiated as previously described) were either untreated or treated with LILRB2 blocker antibody (10µg/mL) for 30mins at room temp in 10% serum-rich RPMI with growth media supplements. After incubation cells were spun down and resuspended in serum-free RPMI with growth media supplements prior to loading. Coverslips were rinsed in MEM and appropriately conditioned macrophages were plated on each coverslip to create three different experimental conditions: control-fc stripes with human macrophages, Nogo-66 or Nogo-Δ20 stripes with human macrophages and Nogo-66 or Nogo-Δ20 stripes with LILRB2 blocker-treated human macrophages. The cultures were stopped with 4% PFA after at least 16hrs. Stripes and macrophages were visualized using goat anti-Fc and rabbit anti-Iba1 and appropriate secondary antibodies, respectively. NogoA-repelled Iba1+ macrophages were counted when at least 50% of the cell body was located in the space between stripes. Receptor specificity of the NogoA-dependent cell-repulsion was confirmed through LILRB2 receptor blockade using a monoclonal blocking antibody. Data distribution was evaluated for Gaussian distribution using the Shapiro-Wilk normality test. Ordinary one way-ANOVA with post-hoc Tukey’s multiple comparisons tests were used for statistical analysis of cell count data using GraphPad Prism 7 (GraphPad).

### Data Availability

snRNAseq dataset available upon request.

## Acknowledgements

The authors wish to acknowledge the technical assistance and contributions of A.L. Chan, A. Grubman, J. Legrand, A.M. Tichy, M. de Souza, J. Homman-Ludiye and C.G. Sobey. This work was supported by grants from the NHMRC (APP108197) to J.A.B. at Monash University, and from the NIH (R35 NS097283) and the Falk Medical Research Trust to S.M.S. at Yale University. A NHMRC Senior Research Fellowship (APP1077677) supports J.A.B. An Australian Postgraduate Award Scholarship supports A.G.B.

The Australian Regenerative Medicine Institute is supported by grants from the State Government of Victoria and the Australian Government.

## Author Contributions

A.G.B., L.T. and J.A.B. conceived and designed the research. A.G.B. designed and performed most of the experiments, analyzed the data and wrote the first draft of the manuscript. J.S., M.L. and M.S. performed the snRNAseq and analysis and wrote transcriptomic relevant sections of the manuscript. A.G.B., J.A.B. and L.T. performed the ET-1-induced stroke surgeries. B.C. performed blood work and flow cytometry. W.K. performed human tissue immunolabeling and assisted with surgeries. T.D.M. assisted with the ASCA2+ astrocytes isolation from mouse pups. S.M.S., N.S., and S.K.N. provided intellectual advice and support. L.T. and J.A.B. supervised the project. J.S., L.T., S.M.S. and J.A.B. were involved in manuscript editing.

## Additional Information

**Supplementary Tables 1-3** are available for download as separate xlsx files.

## Extended Data

**Supplementary Figures 1-9** are available below.

**Supplementary Table 4** is available below.

**Supplementary Table 1.** Specificity scores of individual genes from GM cell-type clusters, WM cell-type clusters, and astrocyte subtypes.

**Supplementary Table 2.** Differentially expressed genes between injured and contralateral nuclei, and between NogoA-positive and NogoA-negative nuclei.

**Supplementary Table 3.** Enriched ontology pathways from astrocyte subtypes, between injured and contralateral nuclei, and between NogoA-positive and NogoA-negative nuclei.

**Supplementary Figure 1.**
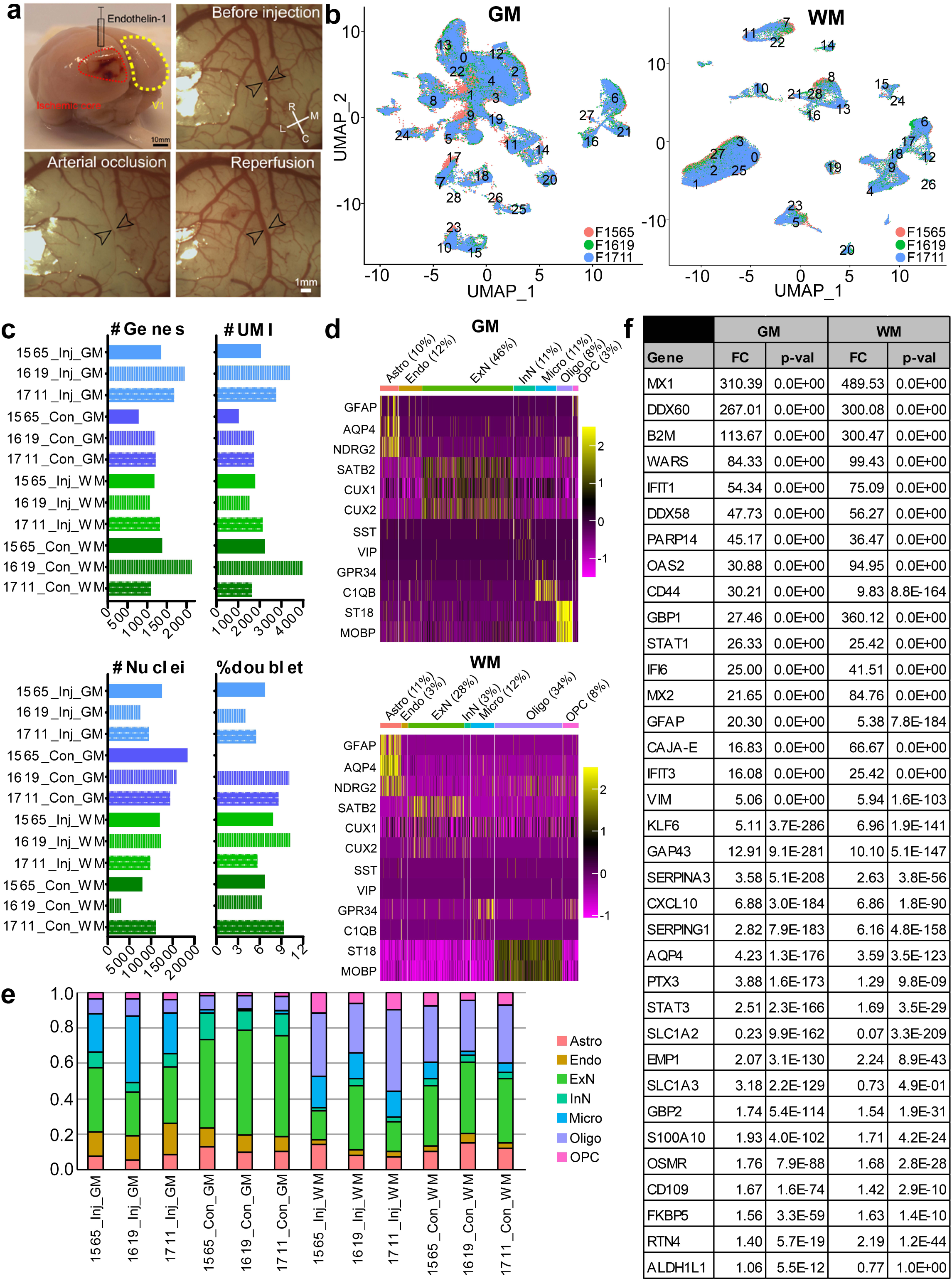
Marmoset ischemic stroke induction with vasoconstrictor and astrocyte identification for transcriptomics. (**a**) Top left; one week post-endothelin-1-induced ischemia of marmoset caudal neocortex (red hatched area ischemic core and yellow hatched area V1 contralateral, used as uninjured control) and pre- (top right)/ post-(bottom images) injection time course highlighting transient ischemia. Arrowheads: occlusion of the posterior cerebral artery. (**b**) UMAP visualization of single astrocyte nuclei, colored by animal. Overlaid numbers correspond to individual clusters. **(c)** Summary of the number of genes, number of unique molecular identifiers (UMIs), number of nuclei, and doublet percentage for each individual sample. Samples are colored by injury and region. **(d)** Heat map colored by single nuclei gene expression of remaining cell-type specific markers; accompanies **Fig. 1c. (e)** Summary of cell-type percentages for each individual sample. Astro = astrocyte, Endo = endothelial cell, ExN = excitatory neurons, InN = inhibitory neurons, Micro = microglia, Oligo = oligodendrocytes, and OPC = oligodendrocyte precursor cells. **(f)** Specific fold change and FDR-adjusted p-values of astrocyte markers and additional genes of interest between injured and contralateral nuclei in GM and WM.

**Supplementary Figure 2.**
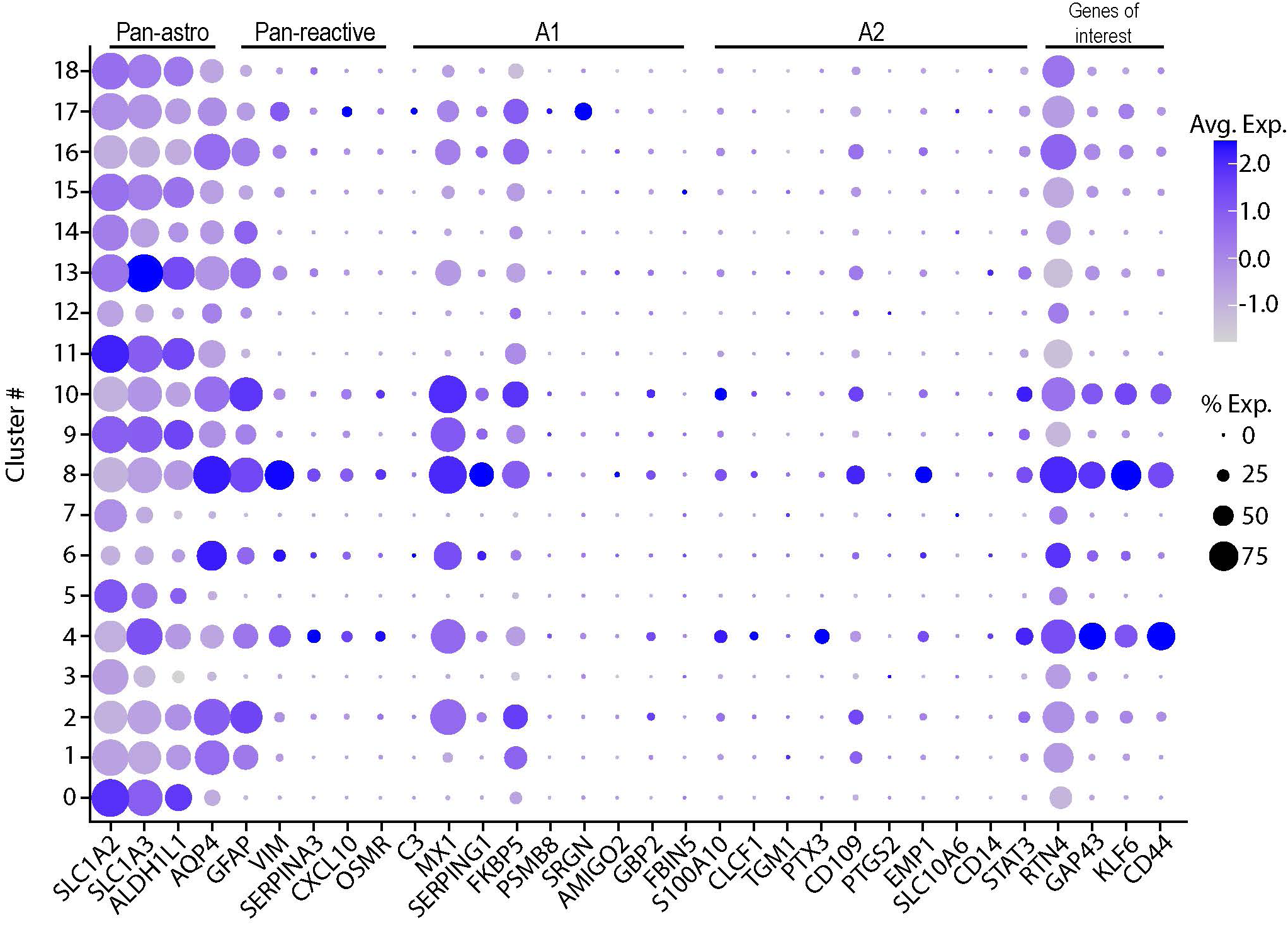
Astrocyte subtype-specific expression of pan-astrocyte and reactive astrocyte markers. The size of the dots indicates the percent of nuclei within that cluster that are expressing the gene at any level, while color indicates average expression from all nuclei within that subtype.

**Supplementary Figure 3.**
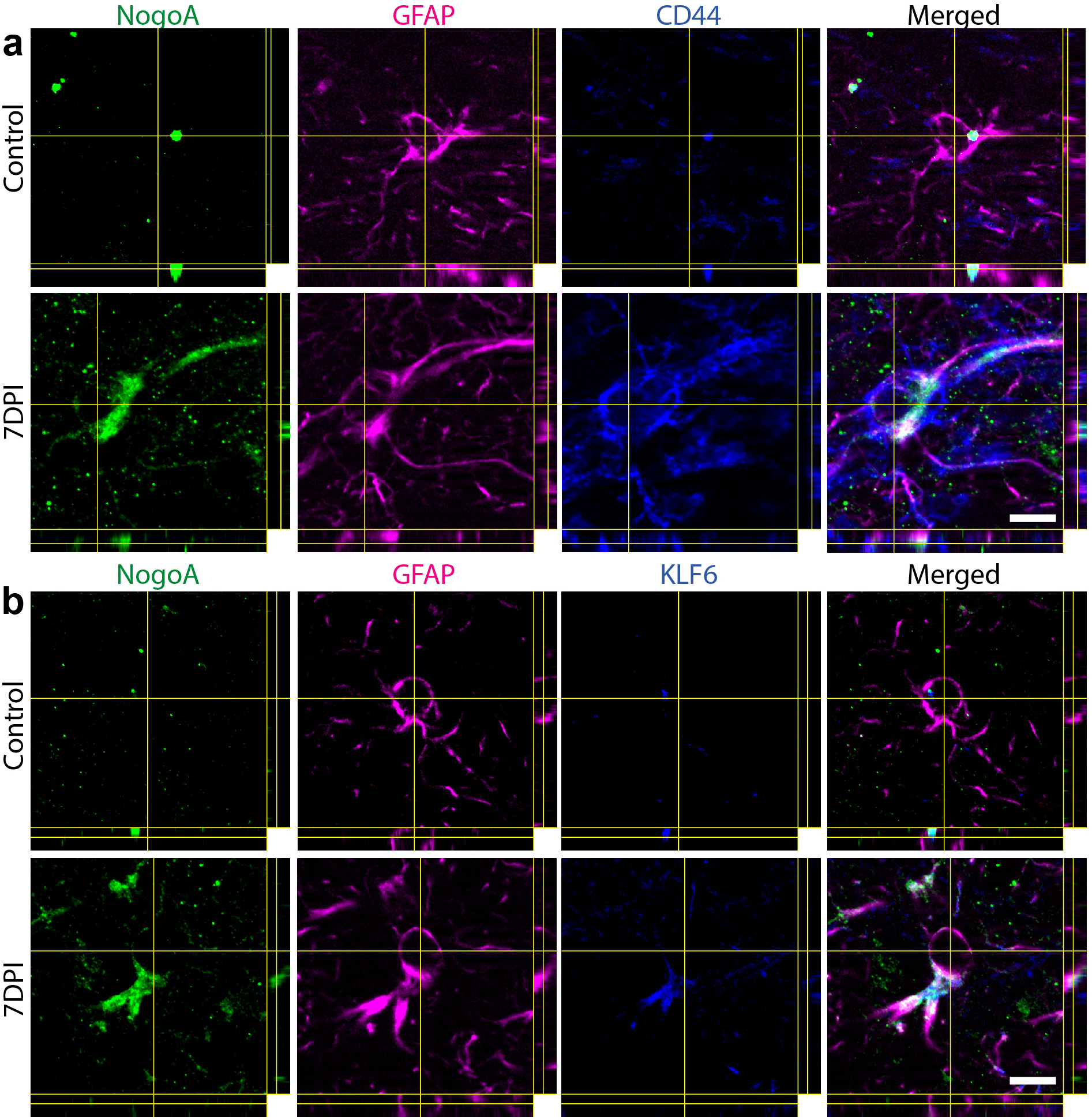
In tissue validation of select markers upregulated concomitantly with NogoA in transcriptomic analysis. (**a**) Validation of CD44 upregulation by immunohistochemistry in marmoset one-week post-stroke, compared to control. (**b**) Validation of KLF6 upregulation by immunohistochemistry in marmoset one-week post-stroke, compared to control. Scale bar: 10µm (**a-b**); CD44: cell-surface glycoprotein-44; KLF6: Kruppel-like factor 6.

**Supplementary Figure 4.**
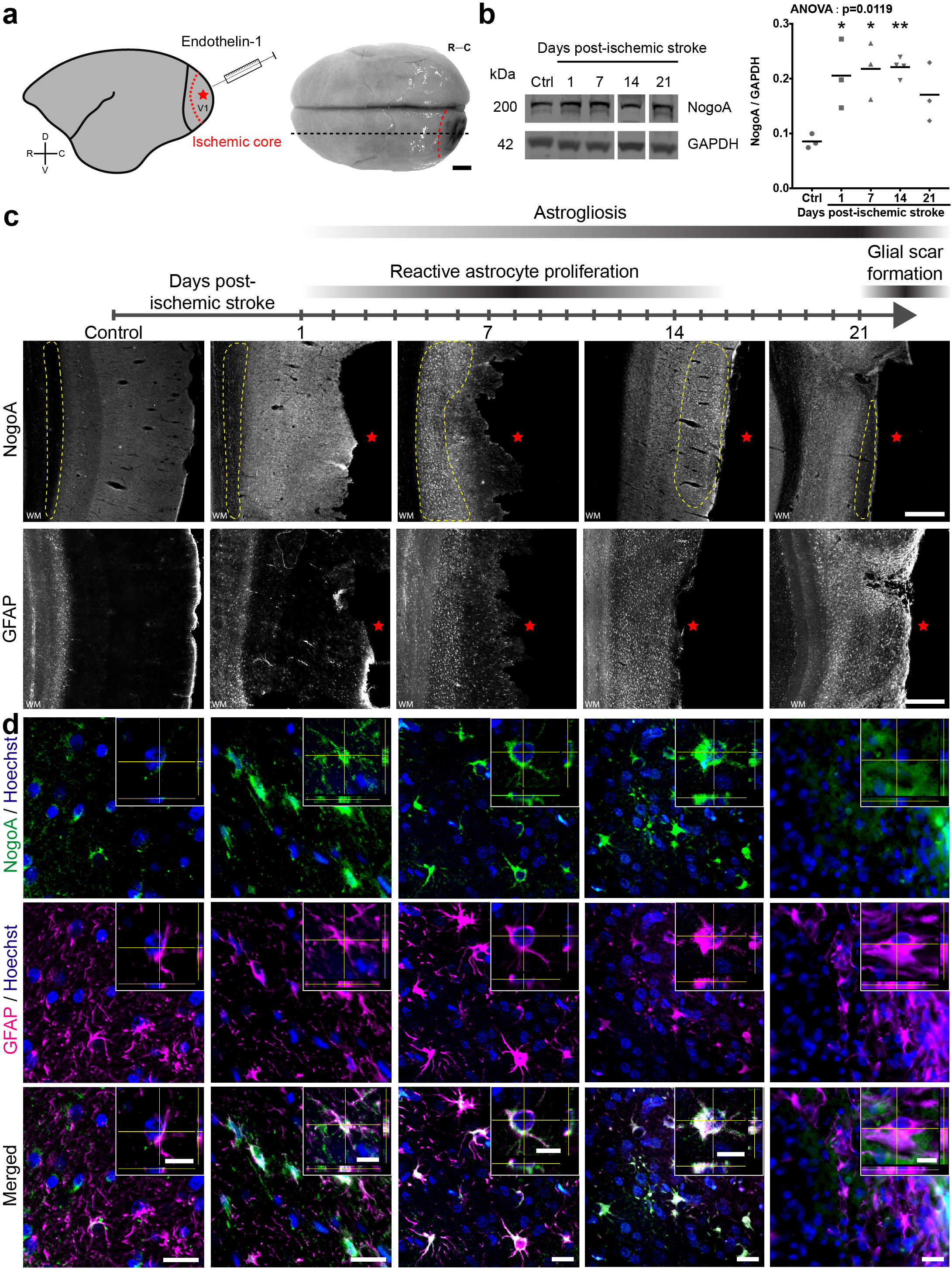
NogoA is upregulated on marmoset GFAP+ astrocytes post-ischemic stroke. (**a**) Schematic and image of marmoset brain depicting ischemic-stroke induction, the region of analysis and lesion size relative to brain size. (**b**) Representative immunoblots for experimental time points analyzed and scatter plots depicting densitometric quantification of NogoA normalized to GAPDH. Each dot represents the mean of 3-4 technical replicates for each marmoset biological replicate analyzed, n=3 per time point, excluding 14 DPI where n=4. (**c**) Schematic of astrocytic pathophysiological time course post-ischemic stroke in primates and representative images showing cellular profile of NogoA and GFAP within marmoset control V1 tissue and ischemic zones at 1, 7, 14 and 21 days post-ischemic stroke (DPI) by DAB immunohistochemistry. (**d**) Representative confocal images and stacks with orthogonal views showing NogoA colocalization with GFAP+ cells within yellow dotted line region of interest from (**c)** in marmoset control V1 tissue and ischemic zones at 1, 7, 14 and 21 DPI. V1: primary visual cortex; V2: secondary visual area; D: dorsal; V: ventral; R: rostral; C: caudal; kDa: kilodalton; NogoA: neurite outgrowth inhibitor A; GAPDH: glyceraldehyde 3-phosphate dehydrogenase; Ctrl: control; statistical test: ordinary one way-ANOVA with post-hoc Dunnett’s multiple comparisons test; *: p<0.05; **: p<0.01; WM: white matter; yellow dotted line: cell population of interest; red star: ischemic core; GFAP: glial fibrillary acidic protein; scale bars: 5mm (**a**), 500μm (**c**), 20μm (**d**: single images), 10μm (**d**: image stacks).

**Supplementary Figure 5.**
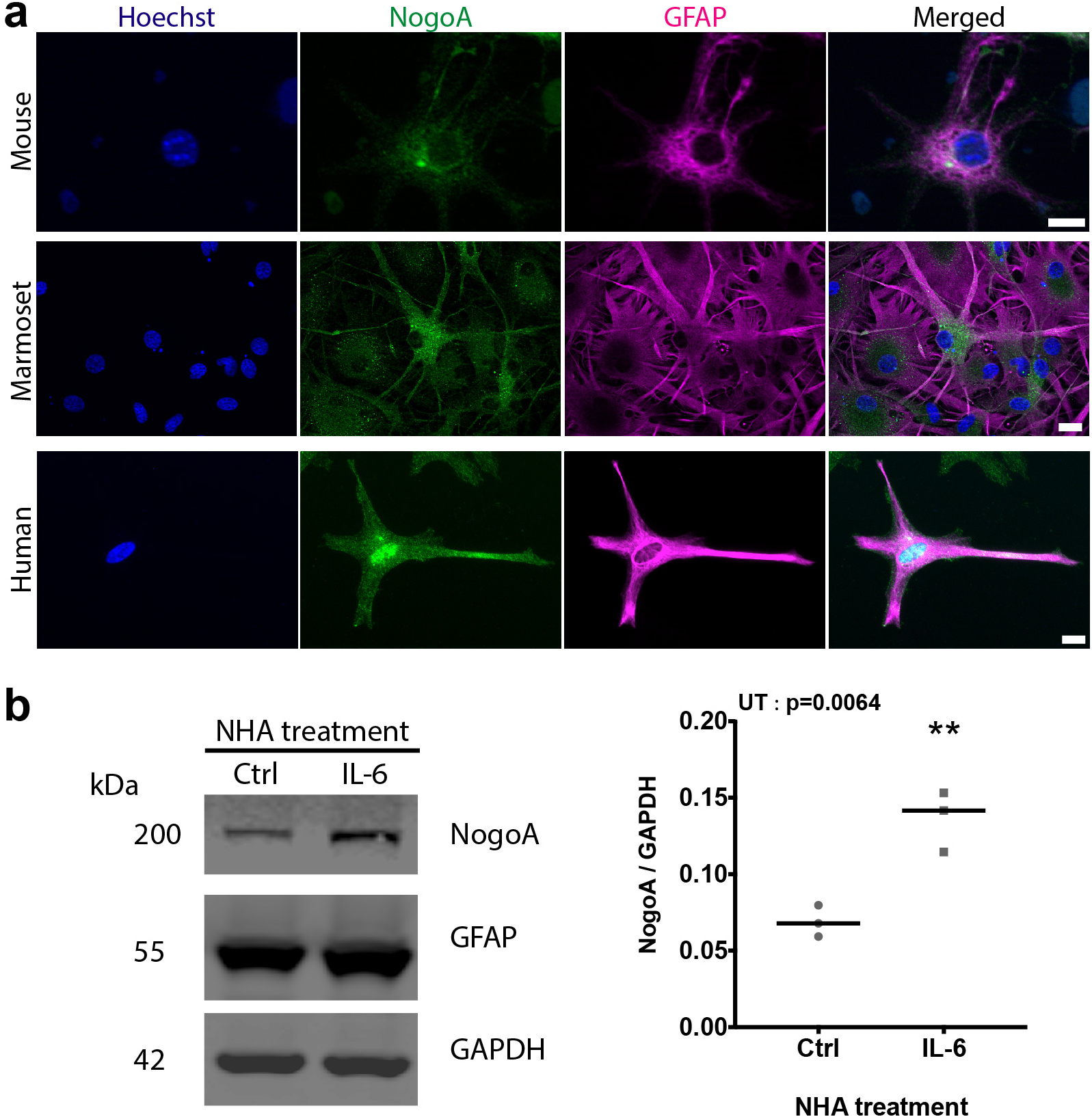
Mouse, marmoset and human astrocytes express NogoA *in vitro*, in the absence of myelin. (**a)** Representative immunofluorescent images showing expression of NogoA on GFAP+ mouse, marmoset and human astrocytes in culture. (**b**) Representative immunoblot and scatter plot depicting densitometric quantification of NogoA normalized to GAPDH. Each dot represents the mean of 3 technical replicates for each cohort of control and IL-6-treated human astrocytes analyzed. GFAP: glial fibrillary acidic protein; NogoA: neurite outgrowth inhibitor A; NHA: normal human astrocyte; scale bars: statistical test: unpaired t-test (UT); **: p<0.01.

**Supplementary Figure 6.**
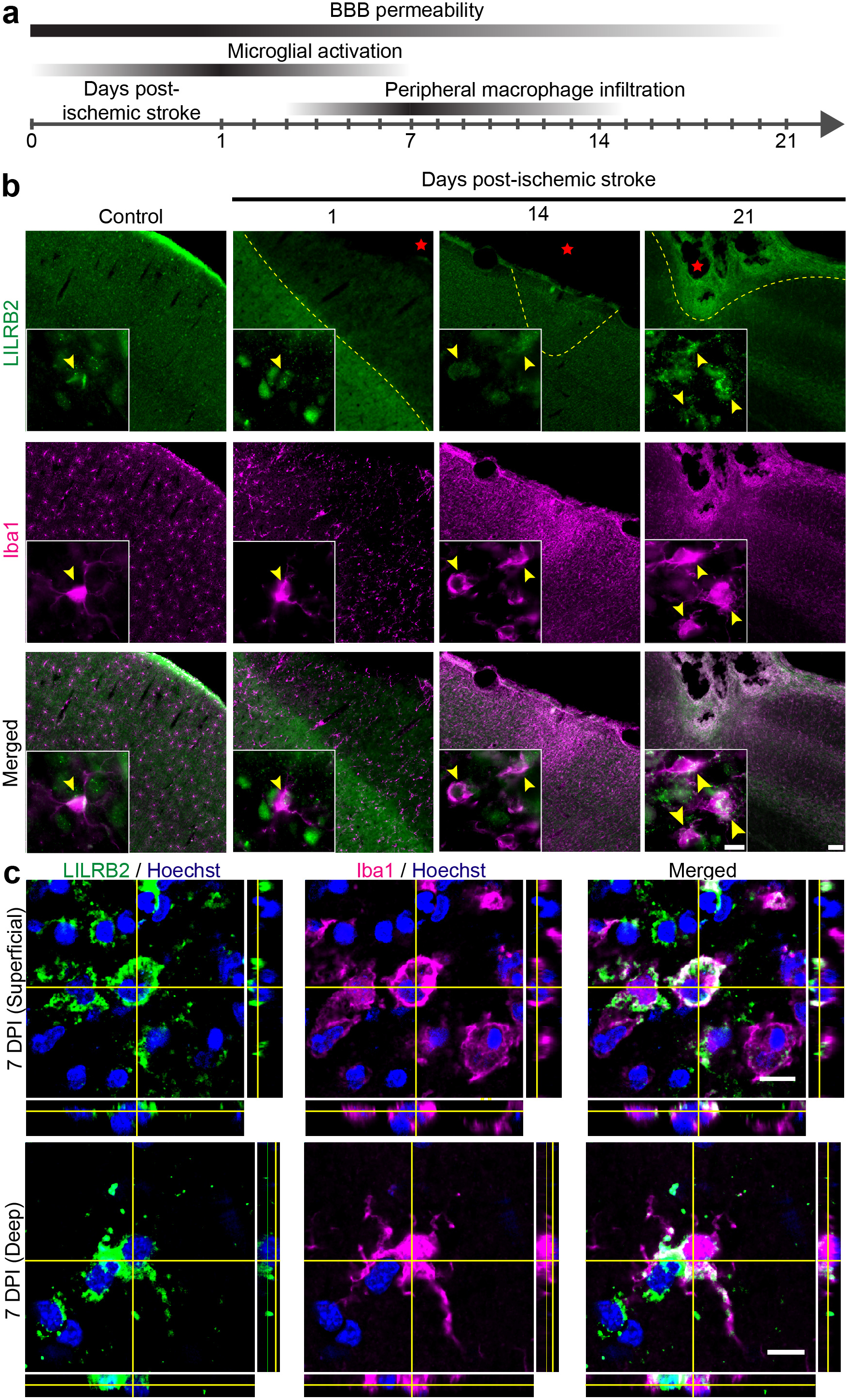
LILRB2+/ Iba1+ macrophages at relevant post-ischemic stroke time points. (**a**) Schematic of macrophagic pathophysiological time course post-ischemic stroke in primates. (**b**) Representative immunofluorescent images showing LILRB2 colocalization with Iba1+ cells in marmoset control V1 tissue and ischemic zones at 1, 14 and 21 DPI. For 7 DPI time point refer to Fig. 4e. Magnified boxes depict double positive cells and morphologies identified within yellow dotted line region of interest in B. (**c**) Representative confocal image stacks with orthogonal views showing LILRB2 colocalization with Iba1+ cells 7 DPI in superficial and deep cortical tissue. LILRB2: leukocyte immunoglobulin-like receptor B2; Iba1: ionized calcium-binding adapter molecule 1; yellow dotted line: cell population of interest; red star: ischemic core; yellow arrowheads: LILRB2+/ Iba1+ macrophages; DPI: days post-ischemic stroke; scale bars: 100μm (**b**: larger box), 10μm (**b**: magnified box & **c**).

**Supplementary Figure 7.**
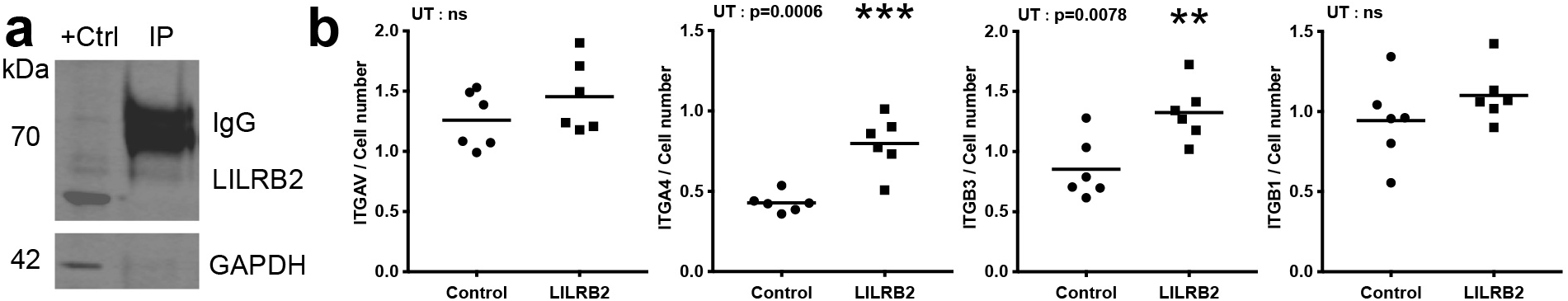
LILRB2 activates integrin signaling. (**a**) Nogo-Δ20 pull down assay of LILRB2 from THP-1 macrophage cell lysate. (**b**) Effect of LILRB2 treatment on THP-1-derived macrophage integrin expression. Scatter plots depict densitometric quantification of protein normalized to cell number. Each dot represents the mean expression of an independent THP-1-derived macrophage cohort analyzed. kDa: kilodalton; +Ctrl: positive control; IP: immunoprecipitation; IgG: immunoglobulin; LILRB2: leukocyte immunoglobulin-like receptor B2; GAPDH: glyceraldehyde 3-phosphate dehydrogenase, ITGAV: integrin subunit alpha V; ITGA4: integrin subunit alpha 4; ITGB3: integrin subunit beta 3; ITGB1: integrin subunit beta 1; statistical test: unpaired T test (UT); **: p<0.01; ***: p<0.001; ns: not significant.

**Supplementary Figure 8.**
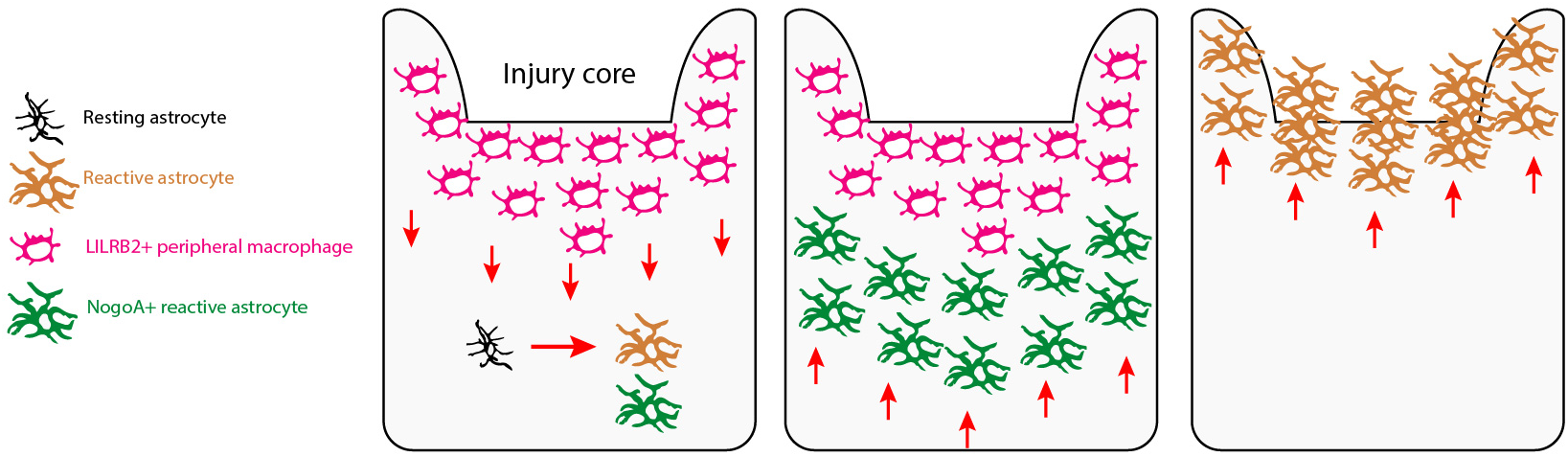
Summary schematic of NogoA-LILRB2 astrocyte-mediated corralling of infiltrating peripheral macrophages following ischemic stroke in primates. LILRB2: leukocyte immunoglobulin-like receptor B2; NogoA: neurite outgrowth inhibitor A.

**Supplementary Figure 9.**
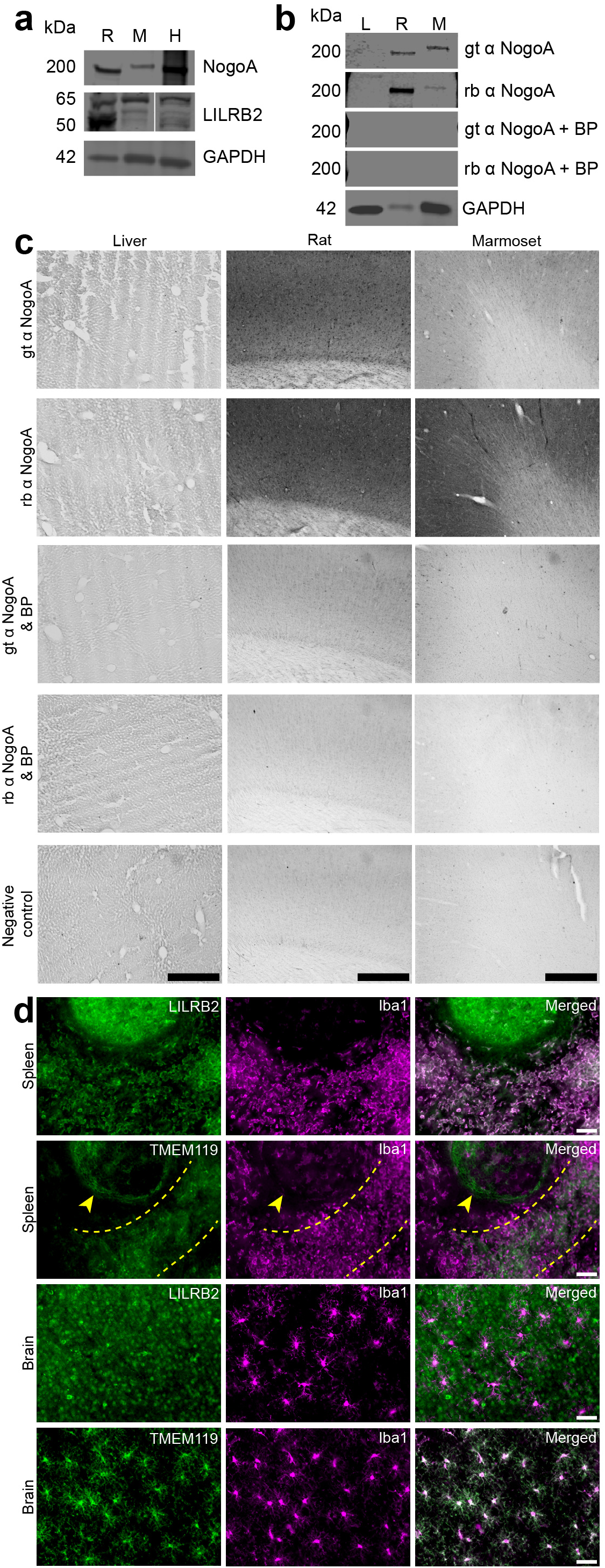
NogoA and LILRB2 antibody characterization in rat, marmoset and human. (**a**) Representative immunoblots of NogoA and LILRB2 in rat (lane 1), marmoset (lane 2) and human (lane 3) cortical tissue. (**b**) Representative immunoblots of two different NogoA antibodies in marmoset liver (lane 1), rat cortical tissue (lane 2) and marmoset cortical tissue (lane 3). (**c**) Representative images show specificity of immunostaining by the two NogoA antibodies used in this study in marmoset liver, rat cortical tissue and marmoset cortical tissue. (**d**) Representative images showing LILRB2 and TMEM119 co-labeled with Iba1 in marmoset spleen and brain. NogoA: neurite outgrowth inhibitor A; LILRB2: leukocyte immunoglobulin-like receptor B2; GAPDH: glyceraldehyde 3-phosphate dehydrogenase; kDa: kilodalton; gt: goat; rb: rabbit; α: anti-; BP: blocking peptide; TMEM119: transmembrane protein 119; Iba1: ionized calcium-binding adapter molecule 1; yellow dotted line: reticular fibers of splenic cord; yellow arrowhead: lymph node capsule; scale bars: 500µm (**c**), 50µm (**d**).

**Supplementary Table 4.**
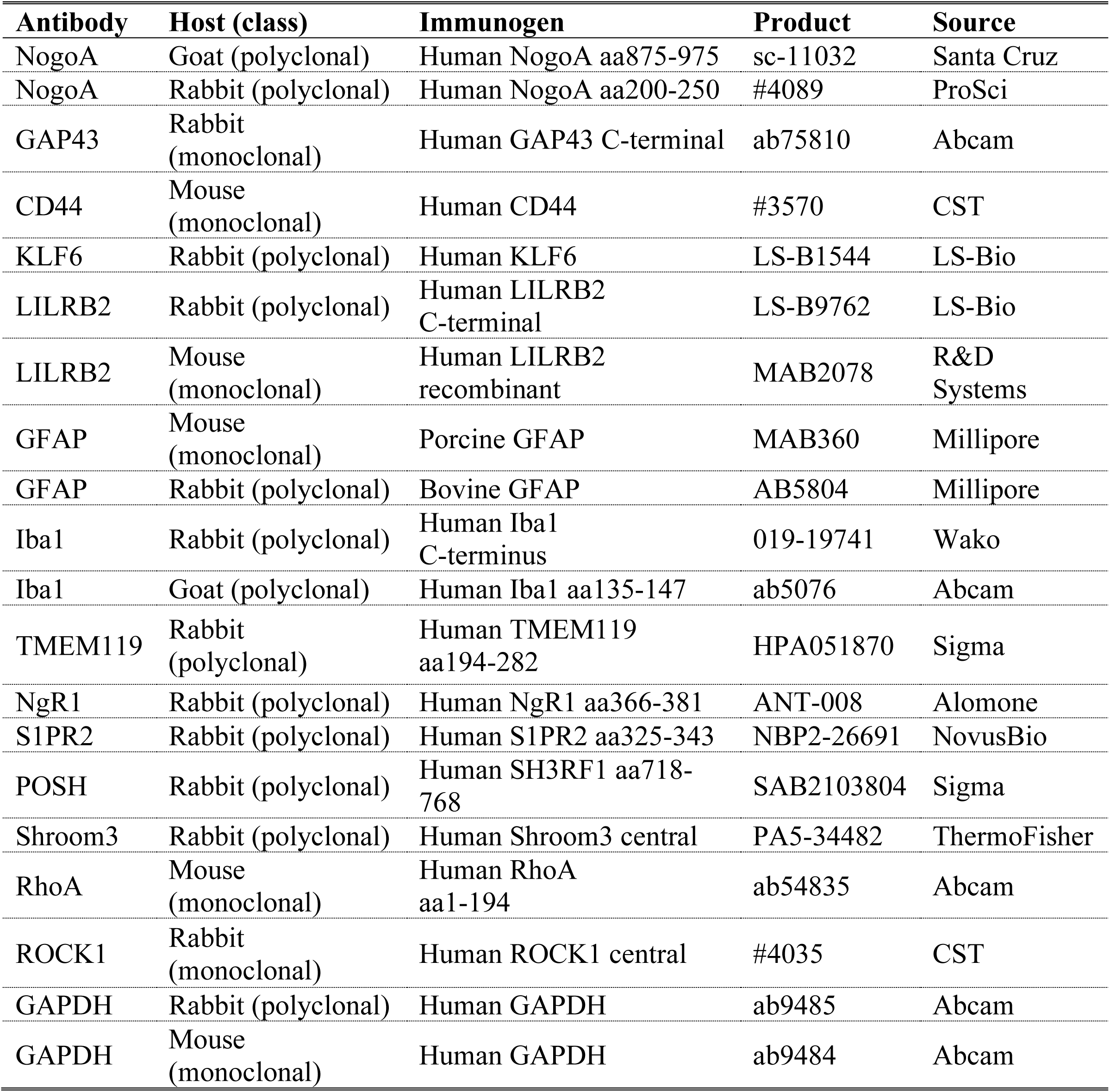
List of primary antibodies used for western blot and immunohistochemistry. NogoA: neurite outgrowth inhibitor A; GAP43: Growth Associated Protein 43; CD44: cell-surface glycoprotein-44; KLF6: Kruppel-like factor 6; LILRB2: leukocyte immunoglobulin-like receptor B2; GFAP: glial fibrillary acidic protein; Iba1: ionized calcium-binding adapter molecule 1; TMEM119: transmembrane protein 119; NgR1: Nogo Receptor 1; S1PR2: Sphingosine-1-Phosphate Receptor 2; POSH: plenty of sarcoma homology domain 3; ROCK1: Rho kinase 1; GAPDH: Glyceraldehyde 3-phosphate dehydrogenase.

